# Digest before Ingest: Early Recruitment of Membrane-bound DNaseX to Phagocytic Cups in Macrophages

**DOI:** 10.64898/2026.02.23.707500

**Authors:** Arghajit Pyne, Vivek Pandey, Subhankar Kundu, Sachie Ikegami, Xuefeng Wang

**Author notes:** Correspondence and requests for materials should be addressed to X.W. equal contribution (Arghajit Pyne, Vivek Pandey).

## Abstract

Macrophages engulf and degrade pathogens and cellular debris through phagocytosis. The degradation process was generally believed to occur only after phagosome internalization and maturation. Here, we report an early DNase activity at the nascent phagocytic cup (PC) prior to its closure. Using a fluorescent DNase sensor, we revealed rapid and ubiquitous DNase activity upon PC formation across various macrophage types. We further identified the responsible enzyme as the membrane-bound DNaseX, which is constitutively recruited to the PC during PC formation. F-actin polymerization was found to correlate with DNase activity in the PC, likely by promoting physical engagement of DNaseX with solid DNA materials. Functionally, we show that macrophages degrade extracellular DNA (eDNA) within bacterial biofilms through direct physical contact, clearing the eDNA structures without internalization. These findings reveal a previously unrecognized DNA degradation mechanism operating at the macrophage membrane, suitable for degrading bulky eDNA materials which cannot be directly internalized by macrophages.

## Introduction

Professional phagocytes, including neutrophils, macrophages, and dendritic cells, are crucial for immune defense and homeostasis maintenance. These cells perform phagocytosis to ingest and break down pathogens, as well as senescent and dead self-cells ^1–4^. During phagocytosis, phagocytes form actin-rich, cup-shaped invaginations of the plasma membrane that envelop target particles ^5^. These phagocytic cups (PCs) are subsequently closed at the cup rims through actin polymerization and myosin-mediated constriction ^6^, leading to the internalization of particles into vacuoles known as phagosomes ^7, 8^. Phagosomes undergo maturation by fusing with endosomes and lysosomes, from which phagosomes acquire various hydrolases such as nucleases, proteases, and lipases ^9–13^. These hydrolases degrade a variety of biomolecules that make up pathogens and self-cells, playing a critical role in pathogen elimination and biomaterial recycling. In particular, among these hydrolases, DNases play a crucial role in breaking down pathogenic and self-cell-generated extracellular DNA (eDNA) ^14–16^, which can cause inflammation ^17^, severe autoimmune diseases ^18, 19^ and organ damage ^20, 21^ if not cleared promptly.

Traditionally, it is believed that DNA degradation during phagocytosis begins only after phagosome-lysosome fusion, during which the phagosome acquires DNase II and becomes acidic, creating optimal conditions for DNase II activity ^22^. However, this “digest-after-ingest” mechanism likely limits macrophages from utilizing phagocytosis to degrade bulky DNA materials, e.g., in neutrophil extracellular traps (NETs) ^23, 24^ or bacterial biofilms ^25^. Both are composed of eDNA structures that exceed the size of phagocytes, and are therefore unlikely to be internalized by macrophages in their entirety. Although DNase I and Caspase-activated DNase ^26, 27^ have been proposed to degrade eDNA materials, these soluble DNases could be rapidly diluted in body fluids and may lack the capacity to achieve the localized, concentrated degradation needed to effectively break down solid eDNA released by pathogens or self-cells.

In this study, we investigated the occurrence of DNase activity during phagocytosis and discovered an unexpected early onset of DNase activity within the PC even before its closure. A fluorescent surface-immobilized nuclease sensor (SNS) ^28, 29^, previously developed in our lab, was adopted to coat microbeads which bait the phagocytosis of macrophages. SNS, essentially as a DNA construct, emits fluorescent signals upon its degradation, faithfully reporting on-site DNase activity at the cell-particle interface. We found that DNase activity is ubiquitously strong within the PC irrespective of the sizes of phagocytic targets (polystyrene microbeads of different sizes) or macrophage types. A series of inhibition and immunostaining results established that the DNase in the PC is glycosylphosphatidylinositol (GPI)-anchored membrane-bound DNaseX, which is also known as DNase I-like 1 (DNase1L1) ^30^. The recruitment of DNaseX to the PC was shown to be constitutive, responding to both microbeads and pathogens (here *E*. coli), regardless of the immunogenic material present on the microbeads. We have also demonstrated that macrophages directly degrade eDNA structures in the *S. Aureus*-formed biofilm by physical contact. The early DNase activity in the PC defines a new interaction paradigm between macrophages and eDNA structures. This “digest-before-ingest” mechanism may greatly empower macrophages to efficiently degrade large-sized eDNA structures in NETs and biofilms using localized and concentrated DNaseX on the cell membrane.

## Results

### Strong DNase activity in the unclosed phagocytic cup (PC)

We developed a platform that induces macrophage phagocytosis and enables the visualization of DNase activity in the PC through fluorescence imaging. As shown in Fig. 1A, polystyrene microbeads were deposited on a glass surface and SNS was coated on both the substrate and microbeads. The SNS construct is an 18 base-paired double-stranded DNA (dsDNA) decorated with a quencher (blackhole quencher 2), an Atto647N dye and a biotin tag facilitating the surface immobilization of the SNS (Fig. 1A) through biotin-avidin interaction. The initially non-fluorescent SNS, if degraded by DNase, leaves a dye freed from quenching on the surface, thereby reporting the local DNase activity by fluorescence on the cell-particle or cell-substrate interfaces. The functionality of SNS on a glass surface was validated by monitoring its fluorescence response to soluble DNase I treatment (fig. S1).

**Fig 1.**
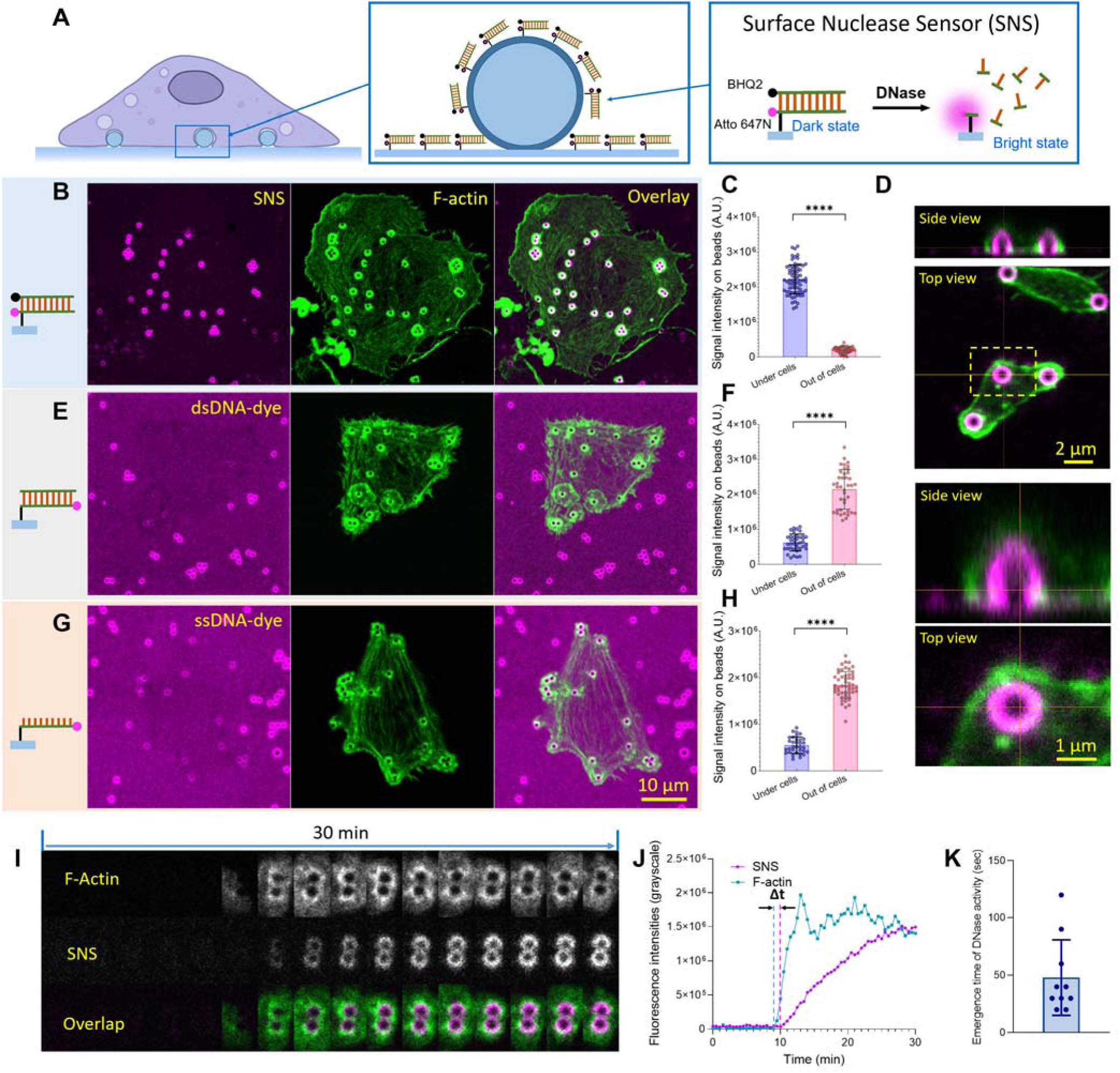
Early onset of DNase activity in the phagocytic cup (PC) prior to its closure. (**A**) Surface-immobilized nuclease sensor (SNS) reports DNase activity in the PCs of macrophages. Microbeads are immobilized on a glass substrate to elicit phagocytosis. The entire surface, including the microbeads, is coated with SNS, a double-stranded DNA (dsDNA) labeled with a quencher-dye pair. SNS becomes fluorescent upon degradation by DNase. (**B**) Human THP-1 macrophages were plated on the DNase-reporting platform, where fluorescence signals appeared in ring patterns, co-localizing with F-actin surrounding the microbeads. (**C**) Comparison of SNS fluorescence intensities on microbeads underneath cell bodies versus those outside of cells. (\*\*\*\**P*<0.0001; Each data point represents one PC; n = 3 experiments; Error bars indicate SD). (**D**) Confocal 3D scanning revealed that the SNS signal was localized on the surface of microbeads. (**E-F**) dsDNA-dye coated on microbeads exhibited a decrease in fluorescence intensities beneath macrophages. (**G-H**) ssDNA-dye coated on microbeads exhibited a decrease in fluorescence intensities beneath macrophages. (**I**) Time-lapse co-imaging of F-actin and SNS signals in the PCs of live macrophages. (**J**) Signal intensity curves of F-actin and SNS on one microbead. The time gap Δt between the starting times of these two signals is defined as the emergence time of DNase activity in the PCs. (**K**) Emergence time of DNase activity in the PCs was statistically estimated to be 48±33 sec after the PC formation indicated by F-actin signal.

Next, human THP-1 macrophages were plated on the microbead-decorated and SNS-coated surface. After incubation for 30 min, the cells were then fixed and F-actin was stained to evaluate the formation of PCs. Fluorescence imaging revealed circular F-actin structures specifically surrounding microbeads, suggesting that macrophages formed PCs in response to the microbeads. Remarkably, strong SNS fluorescence signals, signifying local DNase activity, were observed around the microbeads beneath the macrophages, co-localizing with the circular F-actin signals (Fig. 1B). Note that many SNS-coated microbeads were also present outside of the cell regions, and those microbeads showed no SNS fluorescence signals (Fig. 1C and fig. S2). Moreover, SNS was coated on the entire platform, including both the surfaces of the microbeads and the substrate. However, only the regions of the PCs displayed strong SNS signals, suggesting that SNS signals were highly specific to the PCs formed by the macrophages. To further confirm the specificity and consistency of DNase activity in PCs, large-area imaging was performed on macrophages cultured on fluorescent microbead-decorated and SNS-coated surfaces. The results show that nearly all PCs within macrophages exhibited strong SNS signals, whereas microbeads located outside the cells did not, as shown in figs. S3 and S4.

While SNS signals appear as ring-shaped patterns around the microbeads in two-dimensional imaging, confocal three-dimensional scanning revealed that these signals were actually generated across the entire surface of the microbeads underneath the domes of the PCs, rather than on the substrate regions surrounding the beads (Fig. 1D). To rule out the possibility that the SNS signals in the PCs resulted from dsDNA dissociation due to other factors, such as helicases (albeit unlikely), we further assessed DNase activity in the PCs using dye-labeled dsDNA as a DNase sensor (Fig. 1E). This construct is expected to report DNase activity by fluorescence loss upon degradation, and should not respond to dsDNA dissociation. This strategy of detecting enzymatic degradation through fluorescence loss is similar to the conventional dye-labeled gelatin degradation assay used to evaluate the protease activity of invadosomes ^31^. The result indeed showed that the fluorescence loss was specifically associated with the PCs over the microbeads under THP-1 macrophage cell bodies (Fig. 1E), reinforcing the notion that the dsDNA construct of the SNS was degraded rather than being dissociated within PCs. The average fluorescence loss on microbeads within the PCs is greater than 70%, indicating that the majority of dsDNA was degraded within the PCs (Fig. 1F). Additionally, we tested dye-labeled single-stranded DNA (ssDNA) as a DNase sensor and found that the PCs also caused fluorescence loss of ssDNA on microbeads, suggesting that the DNase in the PCs is capable of degrading ssDNA as well (Fig. 1G and 1H).

We further demonstrated that DNase within the PCs is capable of degrading long dsDNA. As shown in fig. S5, plasmid DNA extracted from *E. coli* was immobilized on a microbead-decorated glass surface through physical adsorption. The plasmid DNA was stained with Sytox green. Macrophages were then plated onto the surface and incubated for 1 h. Fluorescence intensities of Sytox green (DNA stain) associated with microbeads underneath cells were significantly lower than those of microbeads outside the cells, indicating that the plasmid DNA immobilized on the microbeads was degraded by macrophages.

These results revealed strong and specific DNase activity in the PCs of macrophages that degrades both dsDNA and ssDNA, a phenomenon that had not been previously recognized. Because SNS reports DNase activity by fluorescence gain, which has higher specificity and sensitivity than using fluorescence loss, we used SNS as the DNase sensor in the following experiments.

### DNase activity occurs within one minute after the initiation of PC formation

We demonstrated that DNase activity in macrophage PCs occurs prior to cup closure. To further investigate the timing of this DNase activity during PC formation, we performed live-cell imaging of F-actin and SNS signals during phagocytosis. Cells were transfected with Lifeact-GFP, a fluorescent peptide that binds F-actin and enables its visualization in live cells. Due to the high transfection resistance of THP-1 cells, we used mouse RAW 264.7 macrophages as a model for transfection and live imaging.

As shown in Fig. 1I and Video 1, F-actin and SNS signals were imaged in separate fluorescence channels, with the F-actin signal appearing slightly earlier than the SNS signal. Signal intensity analysis in Fig. 1J indicates that the time gap between the onset of these two signals is approximately one minute. Statistical evaluation based on ten PCs determined this time gap to be 48 ± 33 seconds (Fig. 1K and fig. S6), suggesting that DNase activity generally occurs within one minute after the initiation of PC formation. This timing is considerably earlier than the usual duration required for phagosomes to acquire DNase II through phagosome formation, maturation, and fusion with lysosomes, a process that typically takes tens of minutes to hours ^32, 33^.

We also assessed the time required for DNase activity to be detectable in PCs after cell plating. As shown in fig. S7, SNS signals on the microbeads were clearly visible within 10 minutes of cell plating. This result reinforces the notion that macrophages quickly initiate DNase activity in response to phagocytic targets.

Because surface-immobilized microbeads prevent PC closure, they likely prolong PC formation and may alter the observed temporal dynamics of DNase activity within the PC. To determine whether the PC exhibits DNase activity before closure, we prepared SNS-coated free microbeads and fed them to adherent macrophages. The results showed that PCs indeed exhibited DNase activity before cup closure in response to free microbeads (fig. S8), confirming that DNase activity is initiated rapidly during PC formation.

### DNase activity occurred in the PCs of all tested macrophages

After demonstrating DNase activity in the PCs, we proceeded to investigate the ubiquity of this activity across different cell types and phagocytic particle sizes. We tested three types of macrophages: mouse RAW264.7 macrophages, human THP-1 macrophages, and human macrophages derived from monocytes isolated from peripheral blood (fig. S9). Cells of each type were plated on substrates decorated with microbeads of sizes 0.3, 1.1, and 3.0 µm, with both the substrates and microbeads coated with SNS. These cells were incubated for 1 h and fixed for F-actin staining and cell imaging. In all cases, PCs were observed to form on microbeads, indicated by F-actin structures (Fig. 2A-2C). SNS signals were consistently and specifically generated on the microbeads within PCs, irrespective of the bead sizes and cell types. The SNS signal intensities were analyzed and plotted in Fig. 2E-2G. These data suggest that DNase activity is ubiquitous in the PCs.

**Fig. 2.**
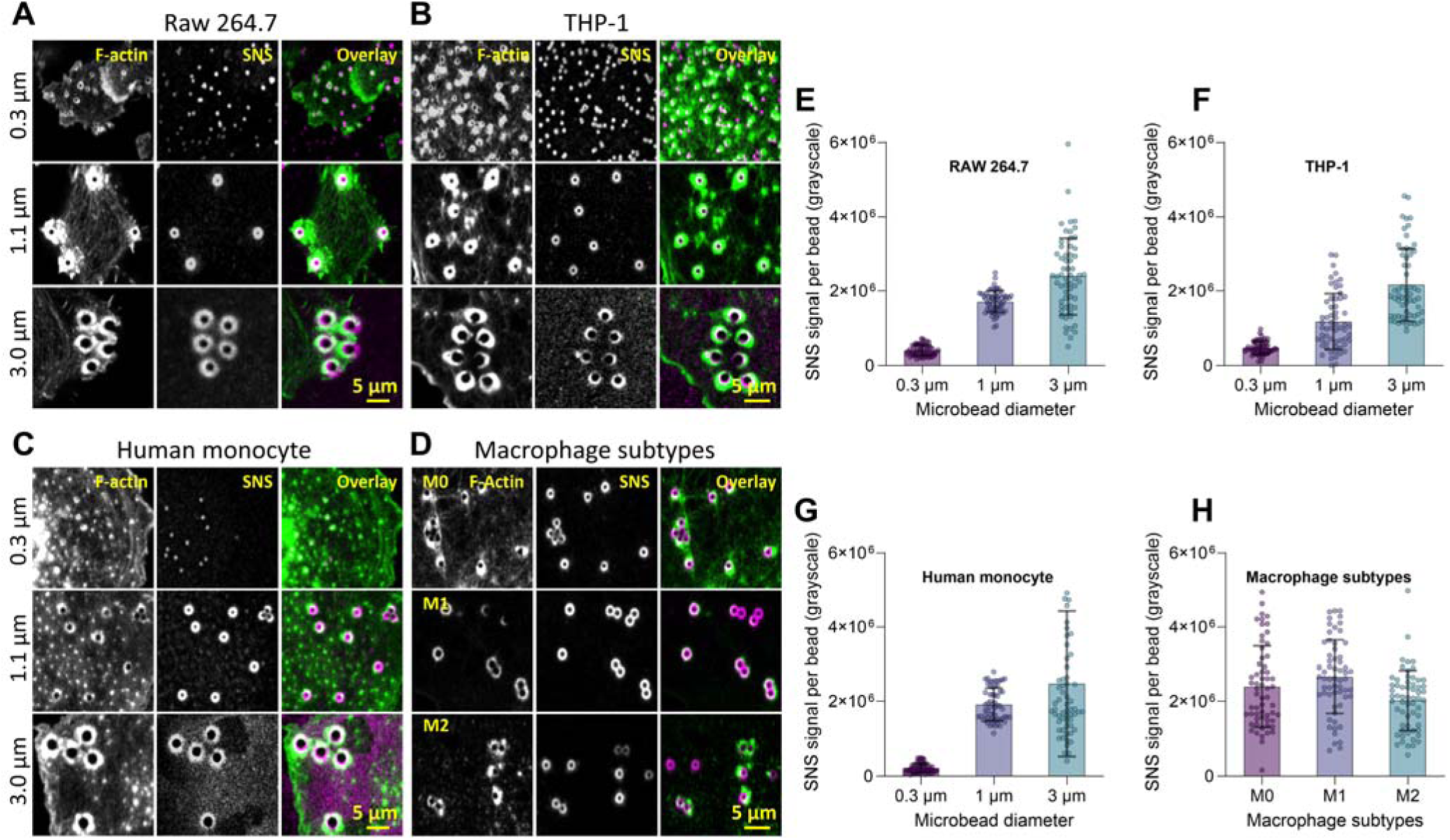
DNase activity is universally present in the PCs of various macrophage types. (**A-C**) DNase activities were consistently observed in the PCs of mouse RAW macrophages, human THP-1 macrophages and human monocyte-derived macrophages. The PCs, marked by F-actin structures, were formed on microbeads of various sizes (0.3, 1.1 and 3.0 µm). (**D**) DNase activity was consistently observed in the PCs of M0, M1, and M2 THP-1 macrophage subtypes. (**E-G**) SNS signal intensities in the PCs of the three types of macrophages. (Each data point represents one PC; n = 3 experiments; Error bars indicate SD). (**H**) SNS signal intensities in the PCs of the three subtypes of THP-1 macrophages. (Each data point represents one PC; n = 3 experiments; Error bars indicate SD)

Dependent on their origins and functions, macrophages generally have three subpopulation: naïve macrophages (referred to as M0 macrophages), pro-inflammatory macrophages (M1 macrophages), and anti-inflammatory macrophages (M2 macrophages) ^34^. Using a protocol introduced in a previous work ^35^, we differentiated THP-1 cells to these three separate subtypes and tested their DNase activities in PCs (see Methods and Materials). The three macrophage subtypes were incubated on 1.1 µm microbead-immobilized, SNS-coated surfaces for 30 minutes. The cell samples were then fixed, and F-actin was stained. The results revealed that all three macrophage subtypes displayed strong DNase activity in the PCs, as indicated by the SNS signals (Fig. 2D). The SNS signal intensities are comparable among the three macrophage subtypes (Fig. 2H). This result suggests that DNase activity in PCs is common among macrophage subtypes.

### Identification of the DNase in the PCs

After observing DNase activity in the PCs, we conducted a series of tests to identify the specific DNase type responsible for this activity. Although a soluble nuclease DNase II ^22^ is known to be recruited to phagosomes, it is unlikely responsible for DNase activity within PCs. This is because DNase II is acquired from lysosomes after the fusion of phagosomes and lysosomes, which occurs during the maturation stage of phagosomes, whereas DNase activity in PCs occurs during the PC formation stage. Additionally, DNase II is an acid nuclease ^36^ that functions in the low pH environment characteristic of mature phagosomes, a condition unavailable in unclosed PCs. Moreover, the SNS signal in the PCs is highly specific and confined to the microbeads, without degrading the SNS on the substrate regions near the microbeads. This suggests that DNase activity in the PCs is not mediated by soluble DNases, which would be expected to diffuse out of the PCs and unable to concentrate the enzymatic activity on the microbeads.

We previously reported that a membrane-bound DNase, DNaseX (also known as DNase I-like 1), is recruited to podosomes of macrophages and invadopodia of cancer cells ^28^. Here, we hypothesize that the DNase in the PCs is also DNaseX. DNaseX is the only known membrane-bound DNase, which is anchored to lipid membrane by a Glycosylphosphatidylinositol (GPI) linker ^37^. We looked up the gene expression of DNaseX (DNASE1L1) and DNase I (DNASE1) in human hematopoietic cells in the website of Expression Atlas (fig. S10). The results revealed that macrophages and monocytes express the highest levels of DNASE1L1 mRNA among all hematopoietic cells, whereas DNASE1 mRNA levels in macrophages and monocytes are generally lower compared to other hematopoietic cells. The high expression level of DNaseX in macrophages suggests a potentially important, yet currently unknown, role of DNaseX in macrophage functions.

Because DNaseX is anchored to the cell membrane by GPI linker, we first tested if the GPI linker is important for the DNase activity in the PCs. We treated the THP-1 cells with Phosphoinositide phospholipase C (PI-PLC), a GPI-cleaving enzyme, and tested the cells on the SNS platform. The result showed that SNS signal intensity within PCs on microbeads significantly decreased upon the PI-PLC treatment, whereas F-actin structures of PCs remained intact (Fig. 3A). When treated with PI-PLC at a concentration of 5 U/ml, the SNS signal intensity in the PCs was nearly reduced to the baseline level (Fig. 3B). This result suggests that the DNase activity in the PCs is associated with a GPI-anchored nuclease, most likely DNaseX.

**Fig. 3.**
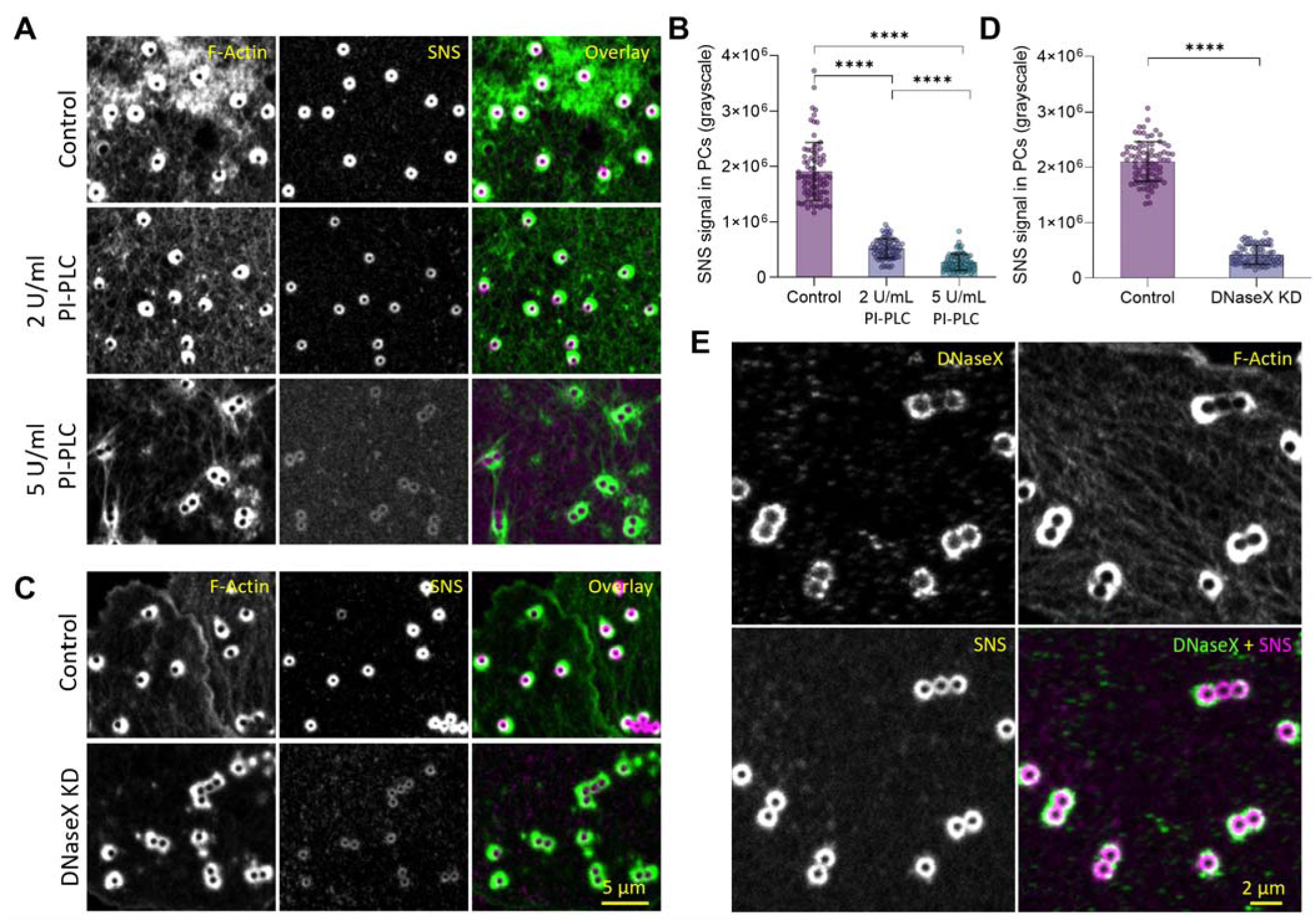
DNase in the PCs was identified as membrane-bound DNaseX. (**A**) F-actin and SNS signals in the PCs of THP-1 macrophages treated with PI-PLC, which cleaves GPI linkers of the putative membrane-bound DNase in the PCs. (**B**) PI-PLC treatment significantly reduced SNS signals in the PCs, indicating marked decreases of DNase activities. (\*\*\*\**P*<0.0001; Each data point represents one PC; n = 3 experiments; Error bars indicate SD). (**C**) F-actin and SNS signals in the PCs of THP-1 macrophages with DNaseX knocked down by siRNA interference. (**D**) DNaseX knockdown significantly reduced SNS signals in the PCs (\*\*\*\**P*<0.0001; Each data point represents one PC; n = 3 experiments; Error bars indicate SD). (**E**) Co-imaging of immunostained DNaseX, F-actin and SNS signals in the PCs of THP-1 macrophages.

Next, we utilized small interfering RNA (siRNA) to reduce DNaseX expression in cells. THP-1 macrophages were treated with siRNA targeting the mRNA of DNaseX, or non-targeting siRNA as a negative control, respectively. While THP-1 macrophages treated with non-targeting siRNA displayed normal SNS signals in the PCs, those treated with siRNA targeting DNaseX showed a significant reduction (p<0.0001) in SNS signal intensities in the PCs (Fig. 3C and 3D). Meanwhile, the F-actin structures of the PCs were formed in both cases, showing no noticeable differences. This result further indicates that DNaseX is the primary DNase responsible for the observed DNA degradation in the PCs.

Finally, we conducted immunostaining using a DNaseX-specific antibody on THP-1 macrophages which were detached by trypsin and cultured on a SNS-coated surface for 1 h. Fluorescence imaging was performed to simultaneously visualize DNaseX, F-actin, and SNS signals. The result shows co-localization among these three signals around the microbeads (Fig. 3E), confirming the presence of DNaseX in the PCs. 3D scan of DNaseX, F-actin and SNS further show that all these signals are on the microbeads beneath the domes of the PCs, not on the glass substrate regions around the microbeads (fig. S11). Based on these lines of evidence, we identified the DNase in the PCs of macrophages as GPI-anchored membrane-bound DNaseX.

### DNaseX is constitutively recruited to PCs without requiring the presence of DNA materials

We established that DNaseX is recruited to the PCs on SNS-coated microbeads. As SNS is essentially dsDNA, an immunogenic substance that could be sensed by DNA receptors such as Toll-like receptor 9 ^38^ or cGAS ^39^, we asked whether the recruitment of DNaseX to PCs is specifically triggered by the presence of DNA on the microbeads. To answer this question, we prepared surfaces without SNS coating and instead decorated them with *E. coli* or microbeads coated with non-DNA biomolecules, including lipopolysaccharide (LPS), rabbit immunoglobulin G (IgG), fibronectin (FN), and poly-L-lysine (PLL). THP-1 macrophages were plated on these surfaces and DNaseX immunostaining along with F-Actin staining were performed (Fig. 4A and fig. S12). Interestingly, the results showed that DNaseX and F-Actin structures were present in the PCs on all particles, irrespective of the biomaterials in the PCs. The confocal images and respective line profile analysis results are shown in the Fig. 4A and 4B. This result suggests that DNaseX recruitment in the phagocytic cups is constitutive, without requiring the presence of DNA materials.

**Fig. 4.**
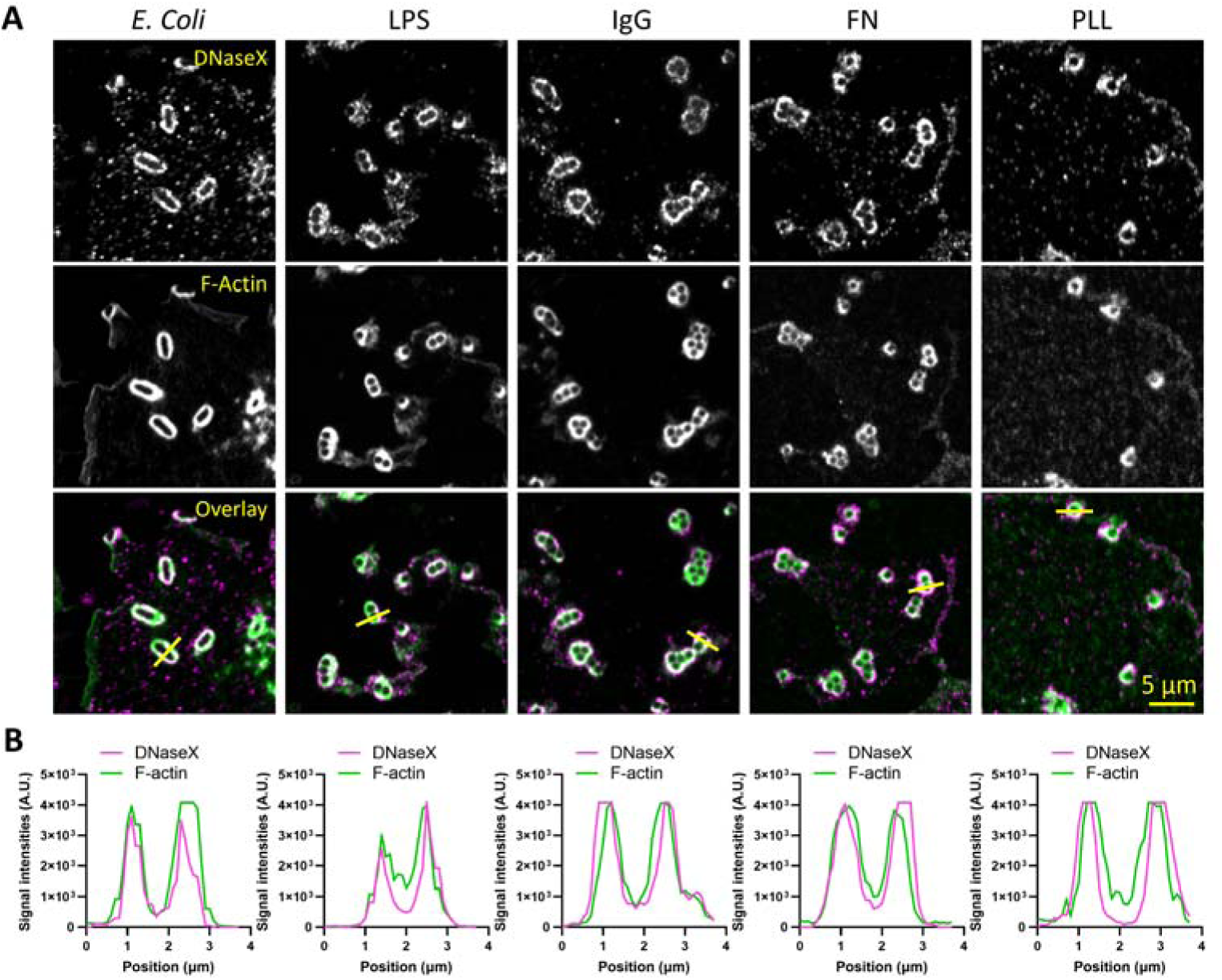
DNaseX is constitutively recruited to the PCs without requiring the presence of DNA materials. (**A**) DNaseX in the PCs in response to surface-immobilized *E. coli*, or microbeads coated with various biomaterials, including lipopolysaccharide (LPS), immunoglobulin G (IgG), fibronectin (FN) and poly-l-lysine (PLL). DNaseX was immunostained with antibodies for imaging. (**B**) Line profiles show the co-localization of DNaseX and F-actin in the PCs on these five surfaces. Yellow lines in (A) mark the locations for the line profile analysis.

### DNaseX in PCs is recruited intracellularly, not from the plasma membrane

As GPI-anchored proteins, DNaseX is synthesized in Endoplasmic Reticulum (ER), processed in Golgi apparatus and transported to the plasma membrane, likely via secretory vesicles. Our results showed that DNase activity is generally specific to the PCs. Here, we investigated the DNaseX distribution on the plasma membrane and in cytoplasm. Because fusion with fluorescent proteins could interfere with DNaseX function by disrupting GPI anchor attachment at the C-terminus or enzymatic activity at the N-terminus, we primarily relied on immunostaining to determine the cellular localization of DNaseX.

We performed immunostaining combined with confocal 3D imaging to examine the subcellular localization of DNaseX in adherent macrophages. To assess membrane association, cells were detached using either ethylenediaminetetraacetic acid (EDTA) or trypsin, which preserve or remove DNaseX from the plasma membrane, respectively. EDTA provides gentle detachment that maintains membrane proteins, whereas trypsin, a protease, degrades membrane-bound proteins during cell detachment. During immunostaining, cell membrane was selectively permeabilized to visualize either total cellular DNaseX or only the membrane-bound fraction. As shown in fig. S13, DNaseX was predominantly detected in submicron-sized plaques on the plasma membrane and in punctate intracellular clusters, likely corresponding to vesicles. DNaseX plaques are not associated with lipid rafts (fig. S14 and Video 2), despite previous reports that GPI-anchored proteins are typically enriched in these membrane domains ^40^.

Since DNaseX is present in both the plasma membrane and cytoplasm, we investigated which region contributes DNaseX to phagocytic cups (PCs) during phagocytosis. To do this, we treated THP-1 macrophages with either EDTA or trypsin, to preserve or remove DNaseX on the plasma membrane, respectively. These cells were then plated on SNS surfaces to observe DNaseX recruitment to the PCs and the corresponding DNase activities. As shown in Fig. 5A, when the cells were treated EDTA, DNaseX existed in both the PCs and submicron-sized plaques which are not associated with phagocytosis. In contrast, when the cells were detached by trypsin, DNaseX was only present at the PCs (Fig. 5B). DNase activities in PCs indicated by SNS signals were normal regardless of cell detachment methods, suggesting that DNaseX in the PCs is not derived from pre-existing membrane clusters but is instead recruited from an intracellular pool of DNaseX within macrophages.

**Fig. 5.**
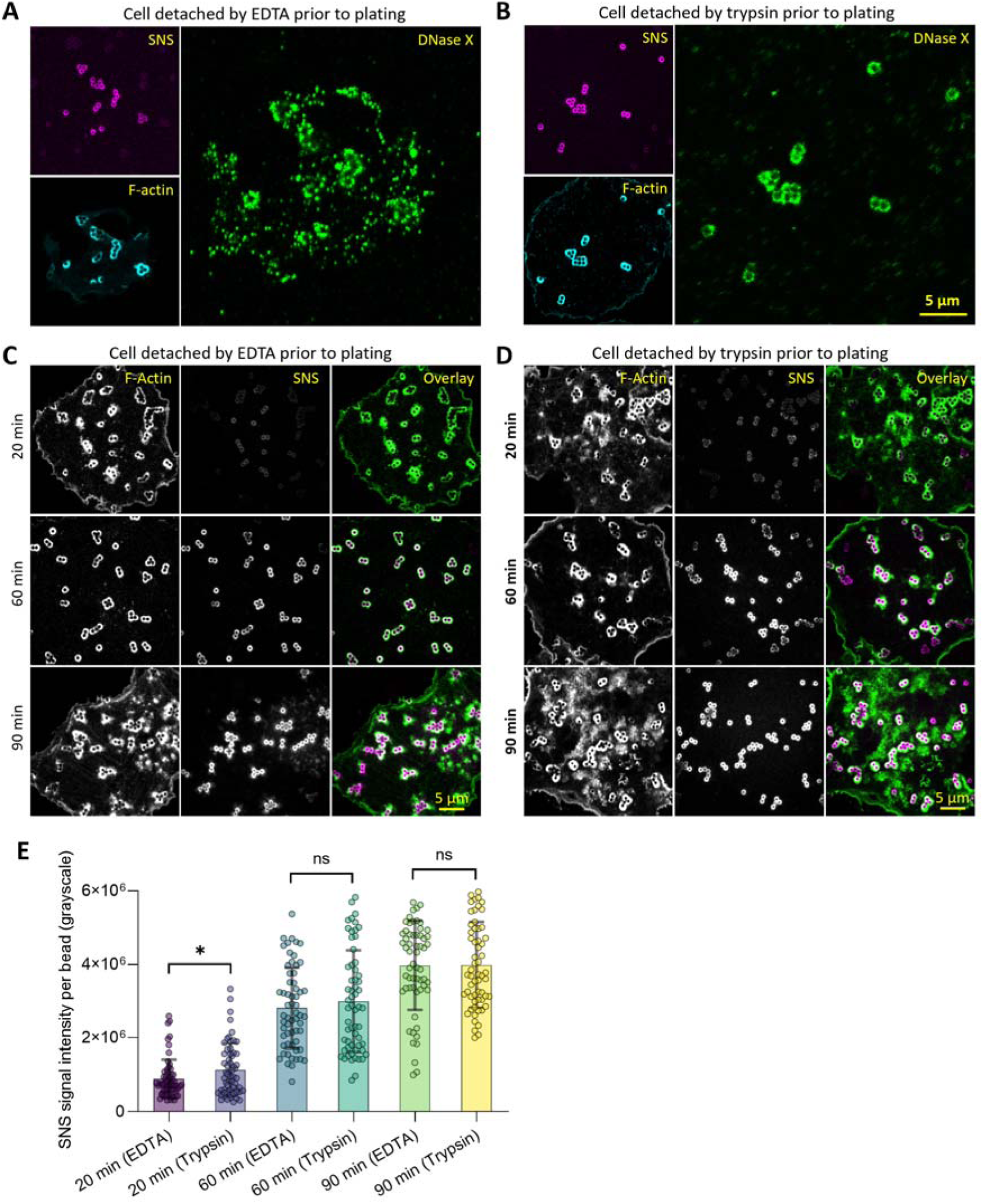
Distribution of DNaseX in the PCs, on the plasma membrane and inside the cell. (**A-B**) SNS signal, F-actin and immunostained DNaseX on the ventral surfaces of adherent THP-1 macrophages, which were detached from a culture flask by either EDTA (A) or trypsin (B) prior to cell plating. The cells were incubated on the SNS surfaces for 1 h. (**C-D**) DNase activities in the PCs indicated by SNS signals with 20 min, 60 min and 90 min incubation times, respectively. The macrophages were detached by either EDTA (C) or trypsin (D). (**E**) SNS signal intensities in PCs. (\**P*<0.05; ns P>0.05; Each data point represents one PC; n = 3 experiments; Error bars indicate SD)

We further compared the time-dependent SNS signal intensities in PCs between EDTA-detached and trypsin-detached macrophages (Fig. 5C and 5D). As shown in Fig. 5E, the differences in DNase activity of PCs between these two conditions were marginal or insignificant at 20, 60, and 90 minutes of cell incubation. This suggests that eliminating DNaseX in the cell membrane by trypsin does not affect DNase activity in the PCs, supporting the idea that DNaseX in the PCs is recruited from the cytoplasm rather than the plasma membrane.

### Actin polymerization is correlated with DNase activity in PCs, but not with DNaseX recruitment

We have consistently observed co-localization of F-actin and DNaseX in phagocytic cups. This observation prompted us to consider the role of F-actin in the DNase activity of DNaseX. Since membrane-bound DNaseX requires direct physical contact with solid DNA materials for degradation, we speculated that F-actin polymerization exerts pressure on the plasma membrane within the PCs, bringing membrane-bound DNaseX into close proximity with solid DNA materials and thereby facilitating the degradation.

To test this hypothesis, we disrupted actin polymerization using 100 µM CK666 or 1 µM cytochalasin D. CK666 inhibits the Arp2/3 complex, thereby preventing actin branching, while cytochalasin D caps the barbed (plus) ends of F-actin filaments, blocking their elongation. We imaged F-actin, DNaseX and SNS signals in macrophages treated with these two reagents or dimethyl sulfoxide (DMSO) as control (Fig. 6A). Both reagents effectively reduced F-actin and SNS signals in the PCs (Fig. 6B and 6C) while DNaseX levels in the PCs were not significantly affected (Fig. 6D). These results indicate that actin polymerization in the PCs is not correlated with DNaseX recruitment, but correlated with its enzymatic activity within the PCs.

**Fig. 6.**
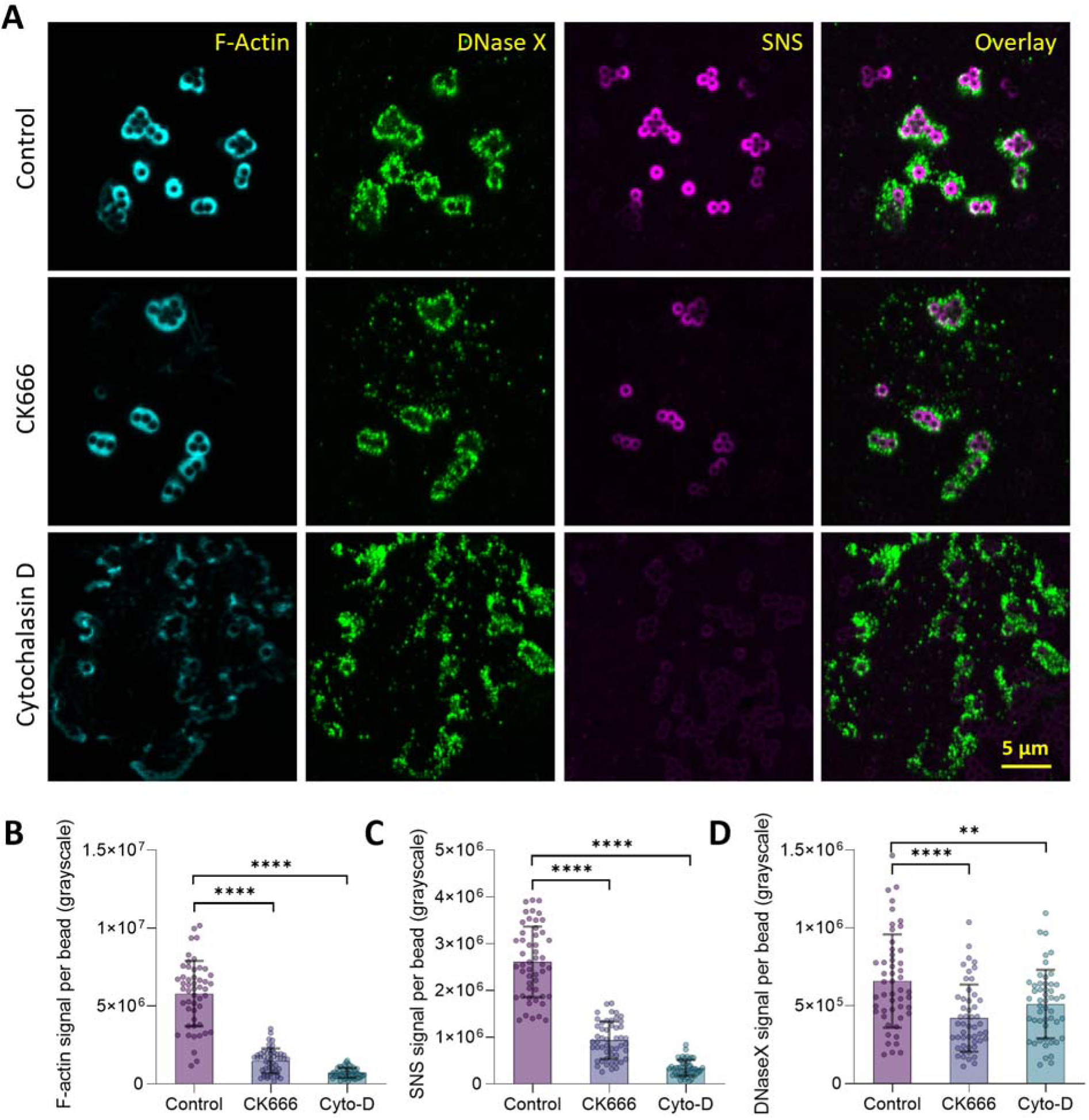
F-actin structure is correlated with DNase activity but not with DNaseX recruitment in the PCs. (**A**) Images of F-actin, DNaseX and SNS signals in the PCs with the macrophages treated with DMSO (control), 100 µM CK666 or 1 µM Cytochalasin D, respectively. (**B-D**) Signal intensities of F-actin, SNS and DNaseX signals in the PCs with the above treatments. (\*\*\*\**P*<0.0001; \*\**P*<0.01; Each data point represents one PC; n = 3 experiments; Error bars indicate SD).

This observation supports the hypothesis that F-actin polymerization generates the protrusive force to push membrane-bound DNaseX to DNA materials for the latter’s degradation. It can also explain why DNaseX in plaques on the plasma membrane generally does not show DNase activity on the SNS-coated surface. This could be due to that DNaseX, as membrane-bound DNase, requires direct physical contact with solid DNA materials on the surface to initiate catalytic degradation, but there is typically a narrow gap, often hundreds of nanometers, between the cell membrane and the underlying substrate ^41^, which effectively blocks the degradative activity of DNaseX plaques.

### Macrophages intensively degrade extracellular DNA in bacterial biofilm by physical contact

A preliminary study was conducted to investigate how macrophages degrade extracellular DNA (eDNA) in bacterial biofilms. *Staphylococcus aureus* (*S. aureus*) is the leading cause of healthcare-associated infections and poses a significant medical and economic burden on society ^42^. These infections are difficult to cure because biofilms formed by *S. aureus* enhance their resistance to antibiotic treatments ^43, 44^, and eDNA is a major structural component of the biofilms ^45, 46^.

We prepared biofilms formed by *S. aureus.* As shown in Fig. 7A, DNA staining indeed revealed filamentous eDNA structures within *S. aureus* colonies cultured on a petri dish for 2-3 days. By a gentle rinse, the bacteria were washed off from the biofilm, RAW macrophages were then plated on the surface and incubated for 30 min. Cells were then fixed for imaging. Fig. 7B shows that eDNA in biofilms was degraded in a pattern highly co-localized with macrophage cell bodies. The boundaries of the eDNA-degraded regions are sharp, with a one micron-scale transition from non-degraded to degraded areas (Fig. 7C), suggesting that RAW macrophages degraded eDNA through direct physical contact rather than via diffusive soluble DNases. Live cell imaging in Fig. 7D and Video 3 further confirm that macrophages degrade eDNA structures within the biofilm, as filamentous eDNAs were progressively severed and dissolved beneath the cell body. A macrophage typically takes 20-30 minutes to degrade eDNA beneath its cell body (Fig. 7E), demonstrating high efficiency in localized eDNA dissolution in bacterial biofilms.

**Fig 7.**
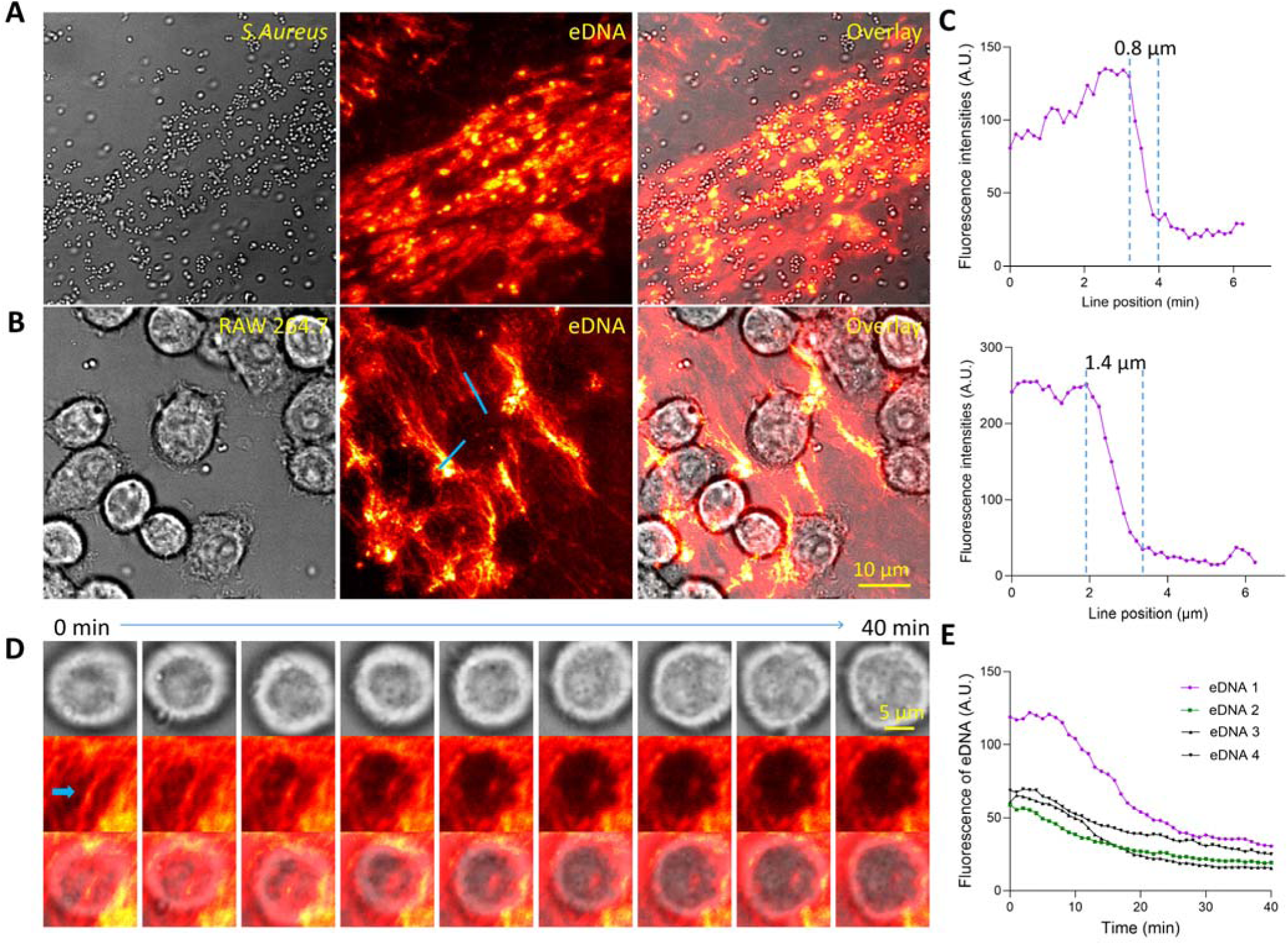
Macrophages degrade eDNA in biofilms by physical contact. (**A**) eDNA structures in *S. Aureus* biofilms. The eDNA was stained with DITO-1. (**B**) RAW macrophages incubated on *S. Aureus* biofilms for 30 min. The filamentous eDNA structures were degraded under cell bodies. (**C**) Line-profile analysis of eDNA degradation regions indicated by blue lines in (B), which are shown to have sharp boundaries with ∼1 µm transition from undegraded region to degraded region, suggesting that the eDNA degradation was mediated by non-diffusive DNase. (**D**) Time-series images of eDNA degradation by a RAW macrophage. Related to Video 3. The eDNA filament (indicated by a blue arrow) became gradually shortened during the degradation. (**E**) Analysis of eDNA degradation by macrophages. The degradation process typically spans 20-30 minutes.

While the results showed that macrophages degrade eDNA in biofilms through physical contact, it has not been confirmed that this degradation is mediated by DNaseX within the PC, as eDNA filaments are not expected to induce the formation of a typical PC structure. An alternative possibility is that DNaseX localized on the cell membrane, including within the plaque-like clusters (fig. S13), comes into direct contact with eDNA and mediates its degradation through physical contact.

## Discussion

Phagocytosis is a crucial process for eliminating pathogens and clearing dead self-cells, playing a vital role in immune defense and homeostasis. Conventionally, PC formation and enzymatic degradation have been viewed as separate stages of phagocytosis. Our findings challenge this view by showing that macrophages recruit membrane-bound DNaseX to PCs at an unexpectedly early stage, coinciding with PC formation. Recruitment of DNaseX to PCs was observed across all macrophage types tested, in response to both pathogens and microbeads coated with various biomaterials. These findings indicate that DNase activity is a constitutive component of PC formation rather than a downstream event. The early and ubiquitous presence of DNaseX in the PCs likely reflects an evolutionarily conserved strategy in macrophages for immune defense. Some pathogens such as Legionella pneumophila have evolved mechanisms to evade lysosomal hydrolases by disrupting phagosome maturation ^47^. By degrading potential extracellular DNA during the PC formation, DNaseX may provide a preemptive assault against pathogenic genetic material, reducing the risk of virulence gene transfer or pathogen-mediated hijacking of host pathways.

In addition to targeting individual micron-sized particles, DNaseX in the PCs may also allow macrophages to efficiently degrade bulky eDNA structures. Both pathogens and decomposing host self-cells release eDNA. Many bacteria integrate eDNA as a major structural material to form bacterial biofilms, which shield bacteria from immune attack of the hosts and antibiotic treatment ^48^. On the other hand, in response to microbial cues and endogenous cytokines, neutrophils can release neutrophil extracellular traps (NETs) ^23, 24^ which are composed of chromatin, to ensnare pathogens and fight infections ^49^. However, excessive accumulation of NETs in organs such as the brain, lungs, heart and kidneys can cause significant organ injuries ^20, 21^, requiring a timely clearance by macrophages. Although soluble DNases such as DNase I ^26^, DNase II ^50^ and Caspase-activated DNase ^27^ have been implicated in DNA clearance, these enzymes have limitations to degrade bulky eDNA structures as soluble DNases can easily be diluted by body fluid and diffuse away from target eDNA. Our discovery of DNaseX in PCs, along with direct imaging of macrophages degrading eDNA in biofilms through physical contact, demonstrates that macrophages utilize membrane-bound DNase to degrade bulky eDNA structures. This process either directly degrades the eDNA structures or process them into smaller fragments suitable for phagocytic uptake. The discovery of membrane-bound DNaseX in PCs provides new insight into how macrophages clear bulky eDNA released by pathogens or damaged self-cells.

Together, the identification of DNaseX in PCs highlights a previously unrecognized mechanism of innate immunity mediated by macrophages. This early, membrane-bound DNase activity equips macrophages with a rapid and spatially focused strategy to neutralize pathogenic or immunogenic eDNA. This discovery opens potential avenues for therapeutic development, such as enhancing DNaseX activity in macrophages to combat biofilm-induced chronic infections and to remove self-DNA that drives inflammation or autoimmune disorders. One limitation of the current study is that the upstream signals responsible for DNaseX recruitment to PCs have not yet been identified. Future work will be needed to define the trafficking pathways of DNaseX recruitment to PCs, and whether its activity can be leveraged to enhance host defense or mitigate DNA-driven inflammatory pathologies.

## Materials and methods

### Synthesis of SNS

SNS is a double-stranded DNA (dsDNA) decorated with a dye, a quencher and a biotin, reporting DNase activity by fluorescence gain. The other two DNase sensors are dye-labeled dsDNA or single-stranded DNA (ssDNA), reporting DNase activity by fluorescence loss. The DNA strands were customized and purchased from Integrated DNA Technologies with sequences and modifications as follows:

**Table 1:**
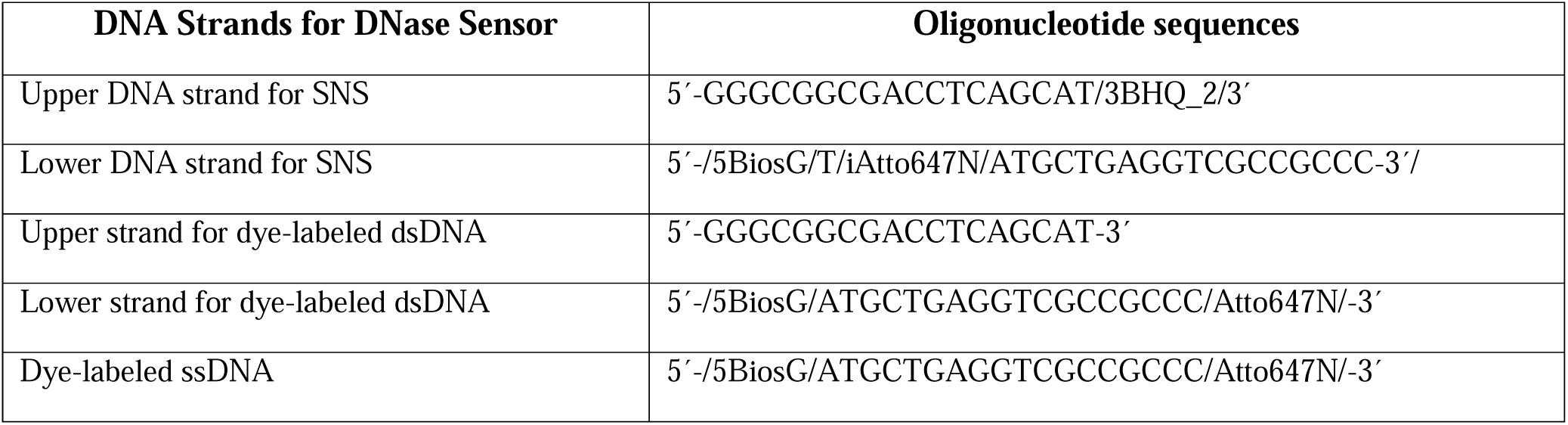
Oligonucleotide sequences for the synthesis of DNase sensors that reports DNase recruitment in the macrophage phagocytic cups.

To prepare dsDNA-based DNase sensors, the respective DNA upper and lower strands were mixed at a molar ratio of 1.1:1.0 and annealed at 90 °C at a final concentration 10 µM. DNase sensors were aliquoted and stored at 4°C for regular use and -20 °C for long-term storage.

### Immobilization of Polystyrene Beads on glass-bottom Petridishes

To study DNase recruitment in macrophage phagocytic cups, polystyrene microbeads were permanently immobilized on a glass-bottom petri dish, serving as the platform for macrophage incubation and imaging. First, a 50 µg/mL poly-L-lysine (PLL; P4707, Sigma-Aldrich) in Milli-Q water was incubated on the glass-bottom petri dish (D-35-14-1.5-N, Cellvis) for 30 minutes at room temperature, followed by three washes with Milli-Q water. Subsequently, polystyrene beads (LB3, LB11 or LB30, Sigma-Aldrich) or fluorescent beads (B0100, ThermoFisher Scientific) were mixed with Milli-Q water at a 1:1000 (v/v) ratio and vortexed for 2.5 minutes. This bead suspension was incubated on the glass-bottom petri dish for 30 minutes at room temperature. The microbeads should be adsorbed on the glass surface by the positively charged PLL coating.

After incubation, the bead solution was carefully drawn by a pipette to minimize liquid residue on the dish. The petri dish was then baked on a hot plate at 120 °C for 1 minute to ensure permanent immobilization of the beads. Following this, the dish was allowed to cool for a few minutes at room temperature and then washed thoroughly with Milli-Q water.

### Coating SNS on the bead immobilized glass surface

To report DNase activity in phagocytic cups, microbead-immobilized glass surface was further coated with the DNase sensor SNS using biotin-avidin interaction. First, a mixture solution of 100 µg/mL BSA-biotin (biotin-conjugated BSA; 29130, Thermo Scientific) and 5 µg/mL fibronectin (FN; 1918-FN, R&D Systems) in PBS (Phosphate-buffered saline, 46-013-CM, Corning) was incubated on the polystyrene bead-immobilized glass-bottom petri-dish for 15 minutes at 4°C. Both BSA-biotin and FN were physically adsorbed on the glass surface. FN was intended to enhance cell adhesion on the surface, and BSA-biotin provides the biotin tag for avidin immobilization, facilitating the subsequent attachment of biotin-tagged SNS.

After incubation, the surface was thoroughly rinsed three times with cold PBS. Next, a solution of 50 µg/mL neutravidin (31000; Thermo Scientific) in PBS was incubated on the petri dish for 15 minutes at 4°C, followed by three washes with cold PBS. Finally, the bead-immobilized surface was incubated with a solution of 0.1 µM SNS in PBS for 15 minutes at 4°C and washed three times with cold PBS. At this stage, the bead-immobilized and SNS-coated glass surface was ready for cell plating and further experiments.

### Coating plasmid DNA on on the bead-immobilized glass surfaces

50 μg/mL PLL in water was added to the well of a microbead-decorated glass-bottom petri-dish and incubated for 20 min. The petri-dish was washed with water three times. Plasmid DNA was extracted from *E. coli* transformed with lifeAct-GFP using high-speed plasmid mini kit (IB47101, IBI Scientific). The immobilization of plasmid DNA is enabled by the electrostatic force as the PLL coating is positively charged and DNA backbone is negatively charged. 50 µg/mL plasmid DNA solution was incubated on a PLL-coated and bead-decorated glass surface for 30 min at the room temperature, and the glass surface was washed with PBS for three times. The DNA was stained with 1 µM Sytox Green and with PBS washed three times. RAW macrophages were plated in the petri-dish and incubated for 1 h in an incubator (5% CO_2_ and 37°C). The sample was imaged directly without cell fixation.

### Coating other biomaterials on the bead-immobilized glass surfaces

To test what biomaterials may elicit the recruitment of DNaseX to phagocytic cups, biomaterials other than DNA materials were coated on bead-immobilized glass-bottom petri dishes. These biomaterials include: 20 μg/ml LPS-EB-Biotin (tlrl-lpsbiot, Invivogen), 20 μg/ml rabbit IgG-Biotin (011-0602, Rockland), 10 μg/ml of fibronectin (37582, Thermo Scientific) in PBS and 50 µg/mL PLL (P4707, Sigma-Aldrich). The biotin-labeled LPS and IgG were immobilized on the surfaces which are pre-coated with BSA-biotin and neutravidin. Fibronectin and PLL were directly coated on the bead-decorated glass petridishes through physical adsorption with 1-hour incubation at 4 °C. These surfaces were washed with cold PBS three times and ready for cell plating.

### Immobilization of E. coli on SNS surfaces

In addition to polystyrene beads, we used *E. coli* (DH5α) as pathogen particles in the phagocytosis study. To immobilize *E. coli*, a glass-bottom Petri dish was coated with 50 µg/ml PLL in milli-Q water for 30 minutes at room temperature, followed by three washes with milli-Q water. The DH5α strain *E. coli* (18265017, ThermoFisher Scientific) or fluorescently labeled *E. coli* (601371, Cayman Chemical) were prepared in PBS and transferred onto the PLL-coated Petri dish, where *E. coli* was incubated for 30 minutes at room temperature. To immobilize the *E. coli* on the PLL-coated surface, 0.5% glutaraldehyde (G6257, Sigma-Aldrich) was added and incubated for 15 minutes. Afterward, the surface was washed three times with milli-Q water and treated with 10 mg/ml sodium borohydride (200050250, Thermo Scientific) to quench any autofluorescence potentially arising from glutaraldehyde. Finally, the surface was washed three times with milli-Q water and ready for macrophage plating.

### Human macrophages derived from peripheral blood mononuclear cells (PBMC)

Human macrophages were acquired by extracting and differentiating monocytes from blood samples. These samples were sourced from leukocyte reduction system (LRS) chambers, a cone-shaped disposable device used in blood transfusion medicine to eliminate white blood cells (leukocytes) from blood products. Consequently, the used LRS chambers contain an enriched level of leukocytes, making them a valuable source for leukocyte extraction.

One used LRS chamber was collected in Hoxworth blood center right after the blood donation (fig S7A). Within one hour after the chamber collection, 5 mL blood sample (Citrate dextrose solution A was used as the anti-coagulant during blood donation) was drained out from the chamber to a 50 mL centrifuge tube. 15 mL PBS buffer (phosphate buffered saline without divalent ions such as Ca^2+^ or Mg^2+^) supplemented with 2 mM EDTA and 2% FBS (fetal bovine serum, 30-2020, ATCC) was added to the blood sample and gently mixed by rotating the tube.

Add 3 mL of density gradient medium (Lymphoprep, Catalog #07801, Stemcell Technologies) to a fresh 15 mL tube. Carefully add 6 mL prepared blood sample on top of the gradient medium. Ensure that a clear separating line forms between the gradient medium and the blood sample. Centrifuge the sample at 400 ×g for 15 minutes using a swing-bucket centrifuge with the brake OFF.

Four layers of substances should be formed in the tube (fig S7B). From top to bottom: The top layer consists of yellow-colored plasma. Below that is a thin and non-transparent layer of PBMCs (peripheral blood mononuclear cells). The third layer is the gradient medium. The bottom layer is red blood cells. Carefully harvest the PBMCs by inserting the pipette directly through the upper plasma layer to second layer where PBMCs are located. Alternatively, the plasma layer can be removed and then collect the PBMCs. The cells are suspended in 2 mL PBS buffer (2 mM EDTA and 2% FBS). PBMCs are now ready for cryogenic storage or monocyte extraction.

### Monocyte extraction from PBMCs

A monocyte isolation kit (Catalog # 19669, Stemcell Technologies) was utilized to extract monocytes via negative selection (Figure S5C). This kit employs paramagnetic beads coated with antibodies that bind to all blood cells except monocytes. Cells captured by the beads are removed from the samples, leaving behind the monocytes. Refer to the detailed protocol provided by the manufacturer of the monocyte isolation kit. The isolated monocytes are cultured in RPMI-1640 medium (30-2001, ATCC) supplemented with 10% FBS and 1× Penicillin-Streptomycin.

### Differentiate monocytes to macrophages

The monocytes were differentiated into macrophages following established protocols: Human monocytes were cultured in a medium supplemented with 50 ng/mL M-CSF (monocyte colony-stimulating factor, 216-MCC, R&D Systems) for four days, and cultured in a medium supplemented with 100 ng/mL M-CSF for another four days The differentiated monocytes became adherent and were harvested using an EDTA-based mild cell-detaching reagent.

### Cell culture

Macrophage cell-lines (RAW264.7 and THP-1) and monocyte-derived human macrophages were cultured and adopted for the study. RAW264.7 and human monocytes were cultured in Dulbecco’s modification of eagle medium (DMEM; 11995-065, gibco) supplemented with 10% fetal bovine serum albumin (FBS; 35-011-CV, Corning) and 1% penicillin/streptomycin (30-002-CI; Corning). THP-1 cells were cultured in RPMI-1640 medium (112-025-101; Quality Biological) supplemented with 0.05 mM 2-mercaptoethanol (1610710; Bio-Rad), 10% FBS and 1% penicillin/streptomycin. We have also checked the culture of THP-1 cells in complete DMEM medium and we did not find any significant difference with respect to cell growth, cell morphology or experimental outcomes. For the macrophage activation, RAW264.7 and THP-1 cells were treated with 0.3 μg/ml phorbol 12-myristate 13-acetate (PMA; P8139, Sigma-Aldrich) for 48 hours. Human monocytes were treated with 100 ng/ml macrophage colony stimulating factor (M-CSF; 216-MCC; R&D Systems) for 48 hours. To generate M1 and M2 macrophages, THP-1 cells activated by the PMA treatment were further treated with 100 ng/ml lipopolysaccharide (L2630, Sigma-Aldrich) or 20 ng/ml recombinant human interleukine-4 (200-04-20UG, PeproTech) for 48 h, respectively. For time-lapse imaging of F-actin dynamics, RAW264.7 cells were transfected with LifeAct-GFP 1-2 days prior to the experiment.

### Cell plating

For experiments, cells were harvested at 80–90% confluency and plated on the microbead/SNS immobilized petri dish surface. According to the needs indicated in the manuscript, cells were detached either by trypsin or by a mild EDTA-based detaching solution (100 ml 10×HBSS + 10 ml 1 M Hepes + 10 ml 7.5% sodium bicarbonate + 2.4 ml 500 mM EDTA + 877.6 ml Water; pH adjusted to 7.4). Cells were incubated in the detaching solution inside a CO_2_ incubator for 5–7 min. Next, cells were gently pipetted off from the culture flask, transferred into a 15-ml centrifuge tube, and centrifuged for 3 min at 300 ×g. The resulting cell pellet was resuspended in complete cell medium and transferred on microbead/SNS immobilized glass surfaces. After incubation in a CO2 incubator for a certain time, the cell samples were ready for follow-up tests. All the experiments were performed using cells dispersed in complete growth medium, except experiments presented in Fig. 4, in which serum-free medium was used instead.

### F-Actin staining and immunostaining

For F-Actin staining and immunostaining, cells were fixed with 4% formaldehyde (28908, Thermo Scientific) solution in PBS for 15 min at room temperature. Next, formaldehyde solution was removed and cells were permeabilized using 0.1% Triton X-100 (BP151-500, Fisher Scientific) solution in PBS for 20 min at room temperature. The cell samples were further blocked with 5 mg/ml BSA (9048-46-8, Fisher Scientific) solution in PBS for 45 min at room temperature. The samples were ready for the follow-up staining.

For F-actin staining, cell samples were treated with a 1:200 dilution of Alexa488-labeled phalloidin (A12379, Invitrogen) and incubated for 30 minutes before imaging. For DNaseX immunostaining, the samples were first incubated with the primary antibody, anti-DNase1L1 (H00001774-MO2; Abnova), diluted 1:200 in PBS, for 1 hour at room temperature. Following this, the samples were incubated with a 1:200 dilution of Alexa405-labeled secondary antibody (A31553, Invitrogen) in PBS, for 1 hour at room temperature. After each incubation step, the samples were washed three times with PBS, with each wash involving a 5-minute incubation.

### Pharmaceutical treatments

To induce GPI-linker cleavage, THP-1 macrophages were incubated with 2 and 5 U/ml of phospholipase C, phosphatidylinositol-specific from *Bacillus cereus* (P5542, Sigma-Aldrich), respectively, for 30 minutes. The cells were then plated and incubated on bead/SNS-coated surfaces for 1 hour prior to fixation and staining. For the inhibition of actin polymerization, Cytochalasin D (C8273, Sigma-Aldrich) and CK666 (182515, EMD Millipore) were added to the cell samples at concentrations of 1 μM and 100 μM, respectively. The cells were incubated with these inhibitors for 1 hour on the microbead/SNS-coated surfaces prior to fixation and staining for imaging.

### siRNA-mediated DNaseX knockdown

To perform the siRNA-mediated DNaseX knockdown, 4 µl of 10 µM siRNA (sc-77165, Santa Cruz) or scrambled siRNA (sc-37007, Santa Cruz) as negative control and 4 µl Lipofectamine RNAiMAX (100014472, Invitrogen) were added to 400 µl OptiMEM (11058-021, gibco) and incubated for 15 min. In parallel, THP-1 cells at 70-80% confluency were detached, centrifuged, and resuspended in serum-free RPMI medium. The transfection cocktail solution was added to the cell suspension and incubated in a 35 mm Petri-dish for 24 h in a CO_2_ incubator. Afterwards, the media of the transfected cells were replaced with complete RPMI medium supplemented with 0.3 μg/ml PMA. After 48 h, the cells were ready for further tests.

### Preparation of bacterial biofilms

*Staphylococcus aureus* (*S. Aureus*) bacterial strain was cultured to prepare bacterial biofilms. 5 mL LB broth was used to resuspend the freeze-dried bacterial powder in the vial (12600, ATCC). 50 µL of resuspended bacteria solution was added to 5 mL LB broth in a tube, which was incubated in a shaker-incubator overnight (∼12 hr) at 37 °C and 200 rpm shaking speed. Afterwards, the cultured bacterial solution was amplified by diluting (1:10 v/v ratio) in the LB broth medium and further incubating inside the shaker-incubator (37 °C and 200 rpm speed) for 4 hr.

For biofilm formation, the amplified *S. Aureus* solution was added to TSB (Tryptic Soy Broth, 41298, Millipore) supplemented with 1% Glucose at 1:6 (v/v) ratio. 400 µL bacterial solution was incubated on glass bottom petridish (D35-14-1.5-N, Cellvis) at room temperature condition for 1-3 days. The biofilm thickness depends on the incubation times, with sporadic thin films formed in 1 day and large and thick films formed in 3 days. The petridish was kept in a pipette box with the bottom chamber filled with pure water to keep the bacterial solution from drying.

After biofilm formation, the surface was thoroughly washed with PBS three times to get rid of any unbound bacteria. The extracellular DNA in the biofilm was stained with DITO-1 (502249754, Aat Bioquest) at 10 µM final concentration and 20 minutes incubation time at room temperature. The sample was subsequently washed with PBS three times. The biofilm was ready for further tests with macrophages.

### Macrophage incubation on bacterial biofilm

To study the interaction between *S. Aureus* biofilm and macrophages, DITO-1-stained biofilm was treated with 0.25% Trypsin-EDTA (CORNING; 25-053-CI) solution and incubated inside the 37 °C incubator for 30 minutes. After incubation, the surface was washed with PBS three times. 200 µL Raw264.7 cell solution was added on the biofilm surface and incubated in a CO_2_ incubator at 37 °C for 1 hr for static imaging. To perform live cell imaging, RAW macrophage solution was plated directly onto the biofilm Petri dish mounted on the microscope. Raw macrophages were detached with mild detaching EDTA solution, centrifuged at 300 rcf for 3 minutes and dispersed in culture medium for live cell imaging.

### Microscopy and image processing

We performed imaging using a confocal microscope (Nikon Eclipse Ti2-E) with a 60× oil immersion objective. For real-time imaging, a lens heater was used to support cell viability. Laser beams at 405, 488 and 640 nm wavelengths were used as the excitation light sources. Image acquisition was conducted using the software (NIS-element) provided with the microscope.

### Quantification and Statistical Analysis

Data were analyzed using Matlab codes developed in our lab. Results were reported as mean ± standard deviation (SD). The results of key experiments were repeated three times and data were pooled together for statistical analysis. The volume of sample analyzed is specified in the figure legends. All the data were analyzed by ImageJ or Matlab codes, and plotted with Graphpad prism. T-tests were conducted using Graphpad prism. For all statistical tests shown: ns: non significant, *p < 0.05, **p < 0.01, ***p < 0.001, ****p < 0.0001.

## ACKNOWLEDGEMENTS

This work was supported by National Institute of General Medical Sciences (R35GM128747). We thank all laboratory members for comments on the study and the manuscript. We are grateful to the quality control team (Stacy Braun, Desmond Taylor, Katheryn Boedecker, Ciara Combs, Sheryl Heeb, Lindsey Marquez, Miriam Michael, Kimberly Molumby, Priddy Taylor, Lindsey O’Bannion, Amy Parker, and Jennifer O’Connor) at Hoxworth blood center for providing the LRS chambers.

## AUTHOR CONTRIBUTIONS

X. W. and A. P. planned the project and designed the experiments. A. P. and V.P. performed experiments and collected data with contribution from S. K. and S. I.. X.W. developed the matlab code for data analysis. A. P. and X.W. analyzed the data. X. W., A. P. and V. P. wrote the manuscript.

## DATA AVAILABILITY

All data supporting the findings of this study are included within the article and its supplementary files. For any further information, inquiries can be directed to the corresponding author, who will respond to and fulfill such requests.

