## Supplementary Figures for "Digest before Ingest: Early Recruitment of Membrane-bound DNaseX to Phagocytic Cups in Macrophages"

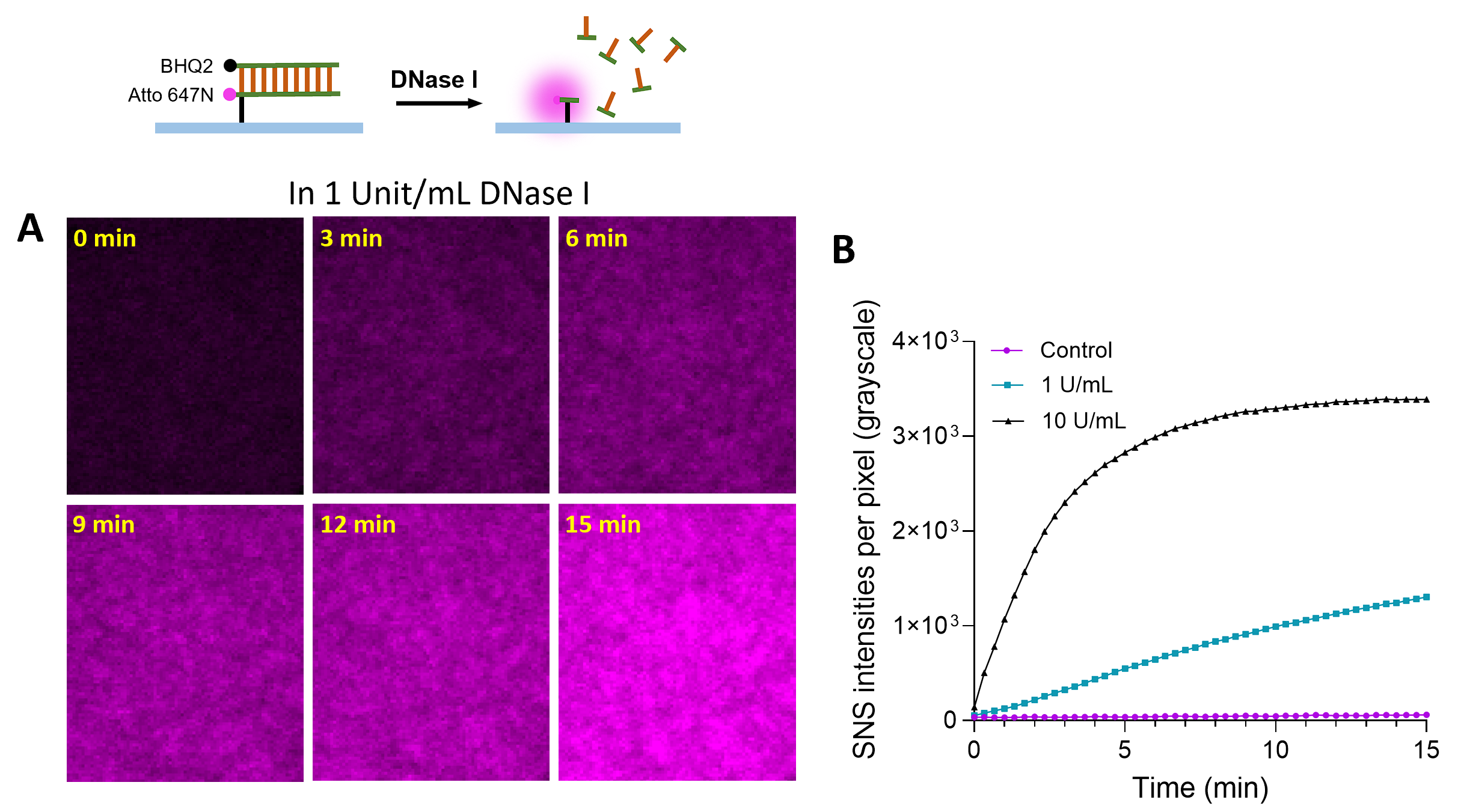

**Fig. S1. SNS immobilized on glass surfaces responds to the treatment of soluble DNase I.**

(**A**) 1 U/mL DNase I (ThermoFisher Scientific, #89836) in Tris buffer (10 mM Tris-HCl, 2.5 mM MgCl₂, 0.5 mM CaCl₂, pH 7.5) was loaded on an SNS-coated glass surface. Fluorescence images were acquired as a time series.

(**B**) SNS response curves to the treatment of DNase I at concentration of 0 (control), 1U/mL and 10 U/mL.

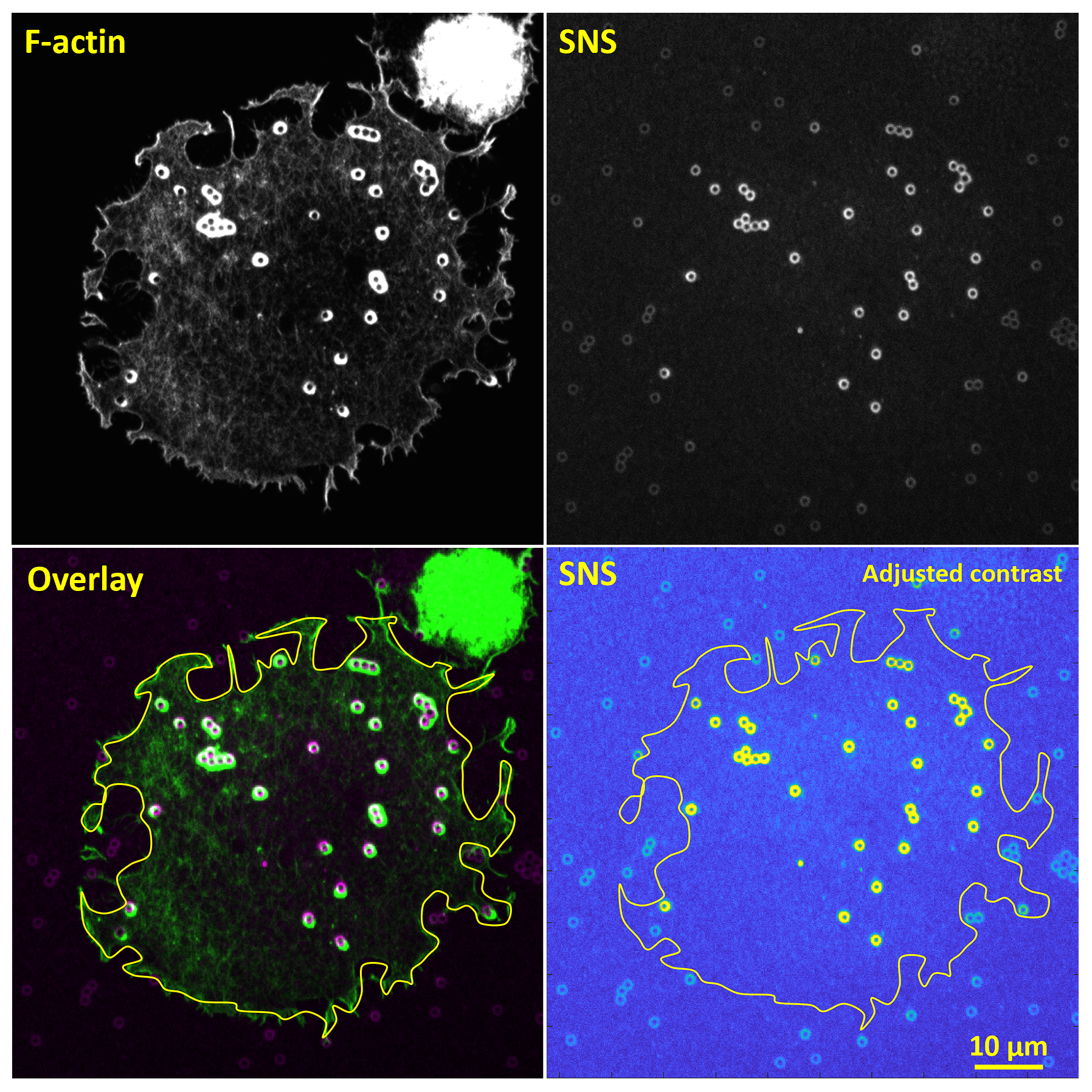

**Fig. S2. SNS signals on microbeads specifically occurred underneath macrophage cell bodies.**

The SNS sensor displays weak fluorescence despite the quencher in the sensor construct for the non-100% quenching efficiency. This low background fluorescence was leveraged to visualize all microbeads on the glass surface by digitally enhancing the image contrast. It was confirmed that the SNS signals, indicated by localized increases in fluorescence, appeared exclusively on the microbeads located beneath the cell bodies of THP-1 macrophages. This observation reinforces the notion that the DNase activity indicated by SNS signals is specific to the cell regions. The yellow contour represents the cell's edge.

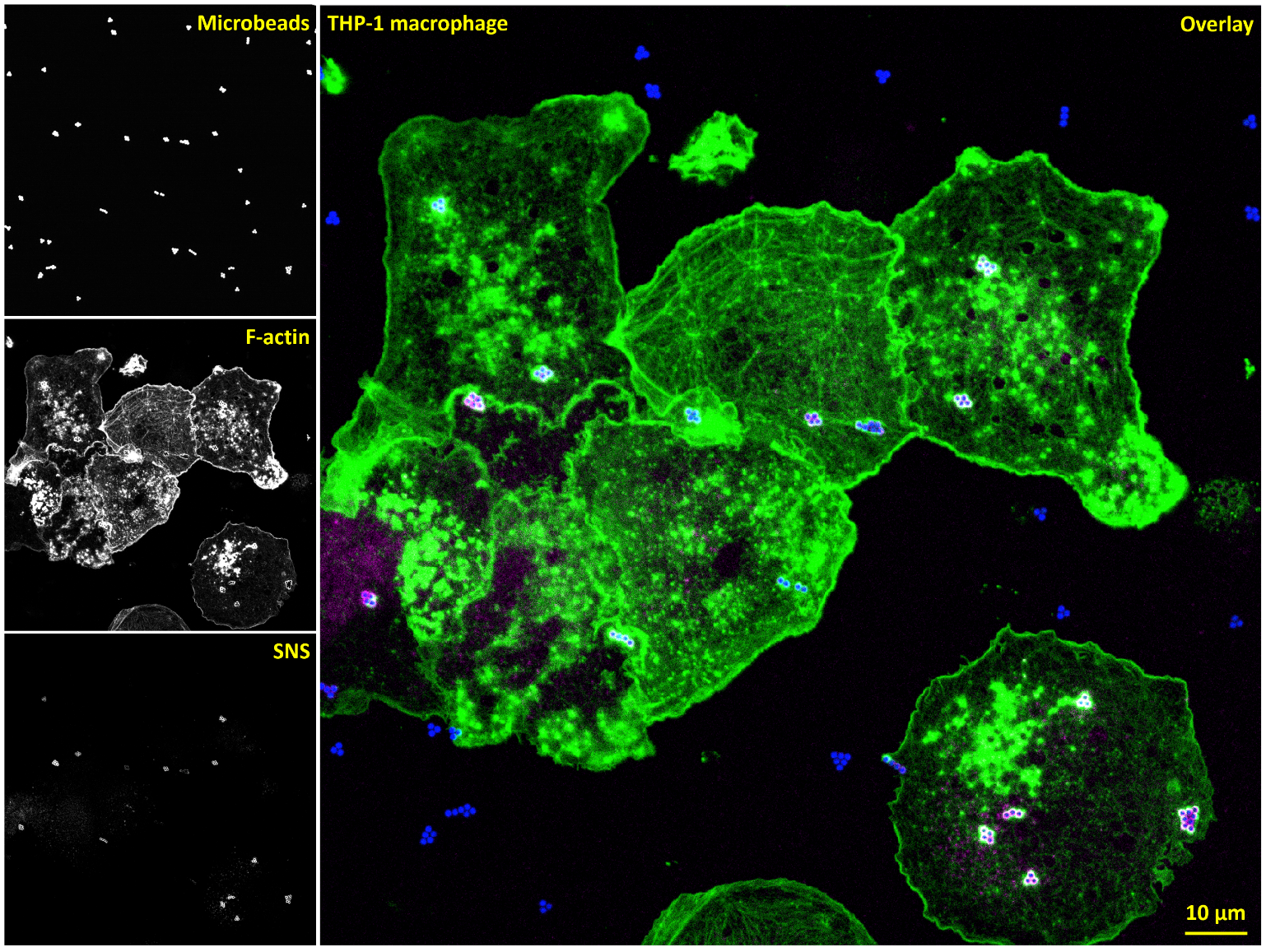

**Fig. S3. Large-area imaging of THP-1 macrophages on an SNS surface.**

Fluorescent beads with an excitation peak at 405 nm were used in SNS experiments to facilitate their localization during fluorescence imaging. Large-area imaging was performed over human THP-1 macrophages plated on an SNS-coated surface, showing microbeads (λ=405 nm, blue), F-actin (λ=488 nm, green), and SNS-reported DNase activity (λ=647 nm, magenta). The macrophages were incubated on the surface for 1 h prior to cell fixation and staining. These images demonstrate that THP-1 macrophages specifically and consistently produce DNase activity in phagocytic cups.

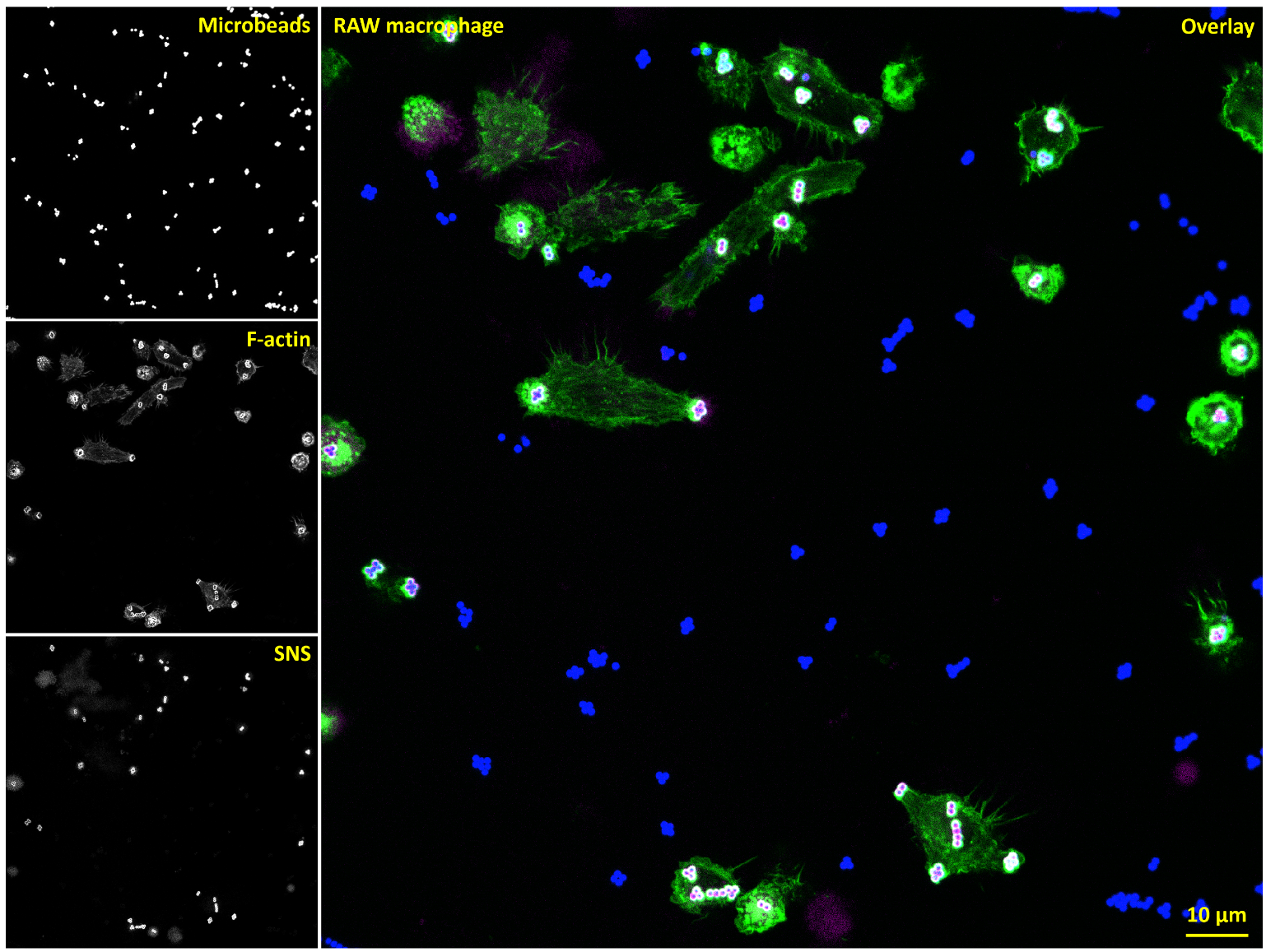

**Fig. S4. Large-area imaging of RAW macrophages on an SNS surface.**

Large-area imaging was performed over mouse RAW 264.7 macrophages plated on an SNS-coated surface, showing microbeads (λ=405 nm, blue), F-actin (λ=488 nm, green), and SNS-reported DNase activity (λ=647 nm, magenta). The macrophages were incubated on the surface for 1 h prior to cell fixation and staining. These images demonstrate that RAW macrophages specifically and consistently produce DNase activity in phagocytic cups.

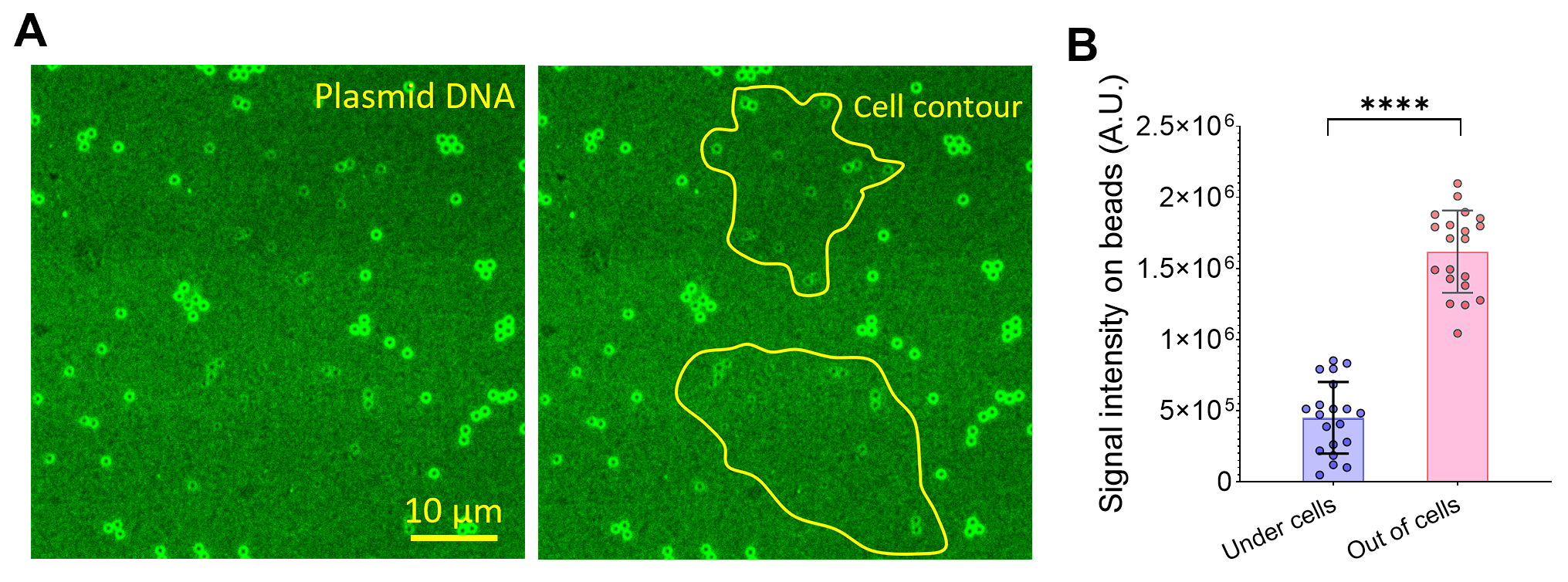

**Fig. S5. The DNase in PCs degrade plasmid DNA.**

(**A**) 50 µg/mL plasmid DNA was immobilized on a microbead-decorated and PLL-coated glass surface. The immobilization of plasmid DNA was enabled by the electrostatic force as the PLL coating is positively charged and DNA backbone is negatively charged. The DNA was stained with Sytox green. RAW macrophages were incubated on the surface for 1 h. The sample was directly imaged without cell fixation or staining. (**B**) Fluorescence intensities of Sytox green on microbeads. The microbeads underneath cells had lower fluorescence intensities than those out of cells, suggesting that PCs degrade plasmid DNA as well.

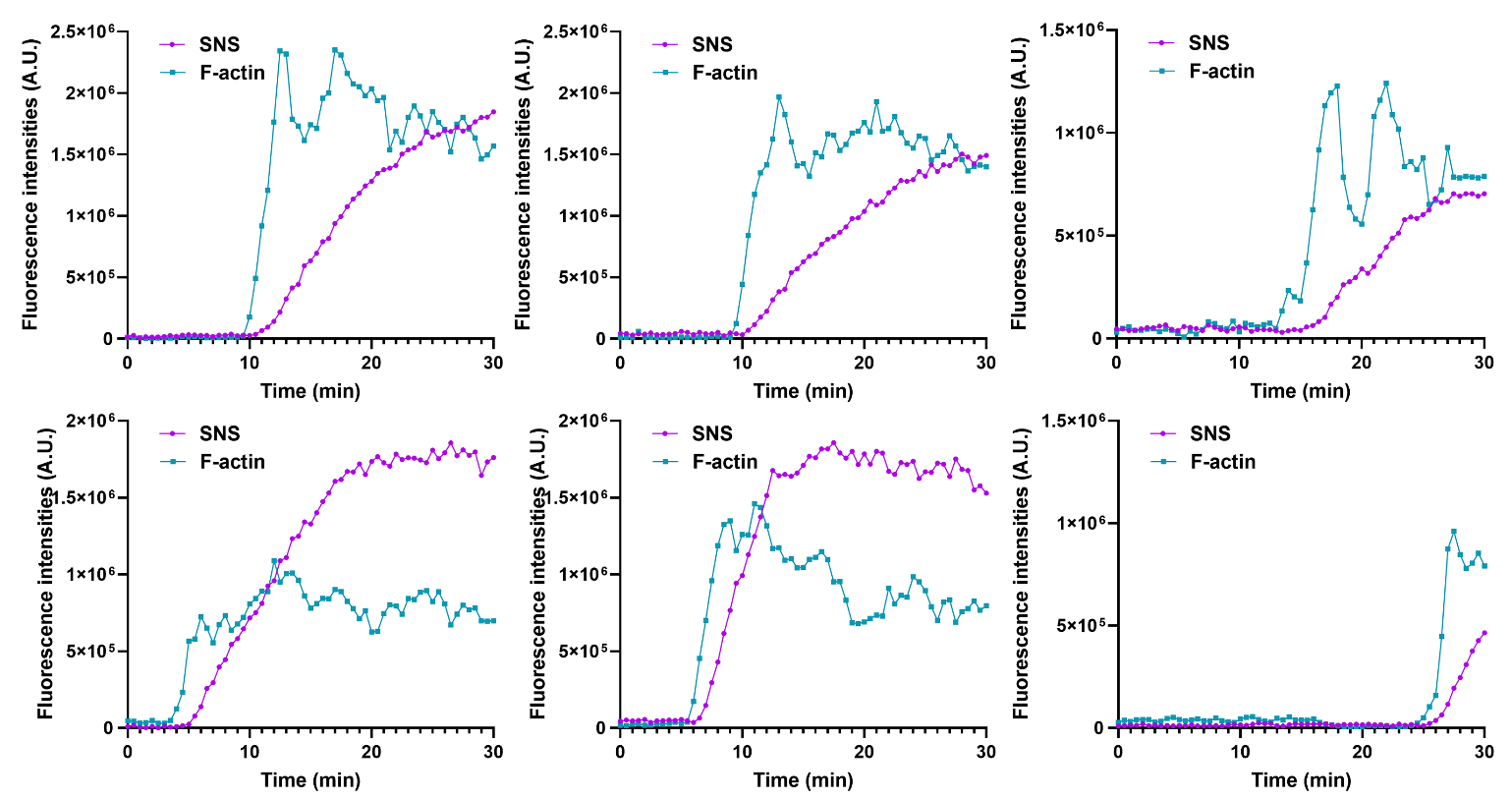

**Fig. S6. F-actin and SNS signal intensities in individual PCs.**

F-actin and SNS signal intensity curves from six additional PCs. Imaging was performed in live macrophages transfected with LifeAct-GFP and plated on SNS-coated surfaces. Related to Video 1.

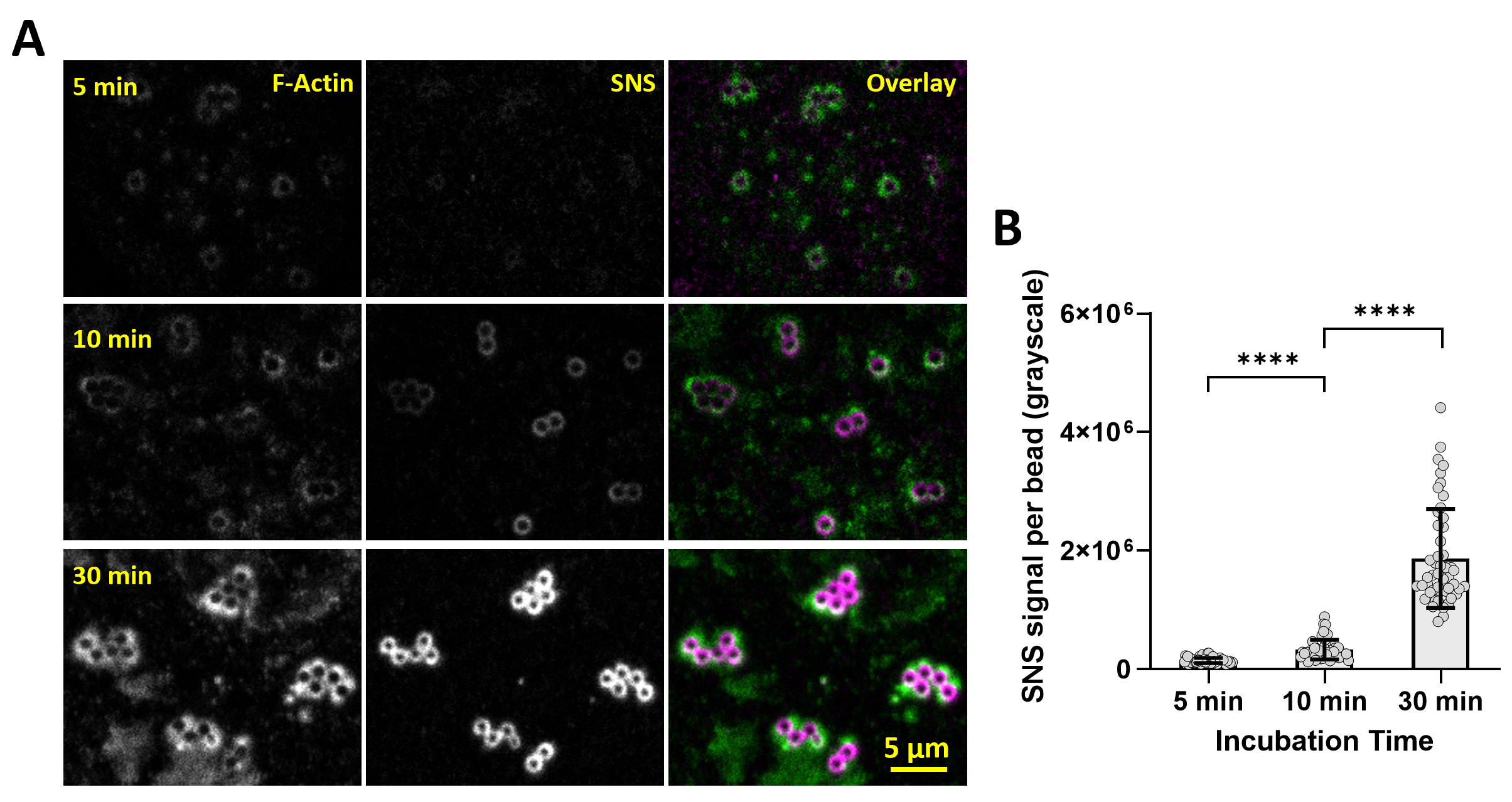

**Fig. S7. SNS signal intensities in the PCs after cell plating for 5 min, 10 min and 30 min respectively.**

(**A**) Co-imaging of F-actin and SNS in the PCs of THP-1 macrophages with 5 min, 10 min and 30 min cell incubation times.

(**B**) Quantification of the SNS signal intensities. Each data point represents the total fluorescent intensity of the SNS signal on a microbead. *****P*<0.0001. Each data point represents the SNS signal in one PC.

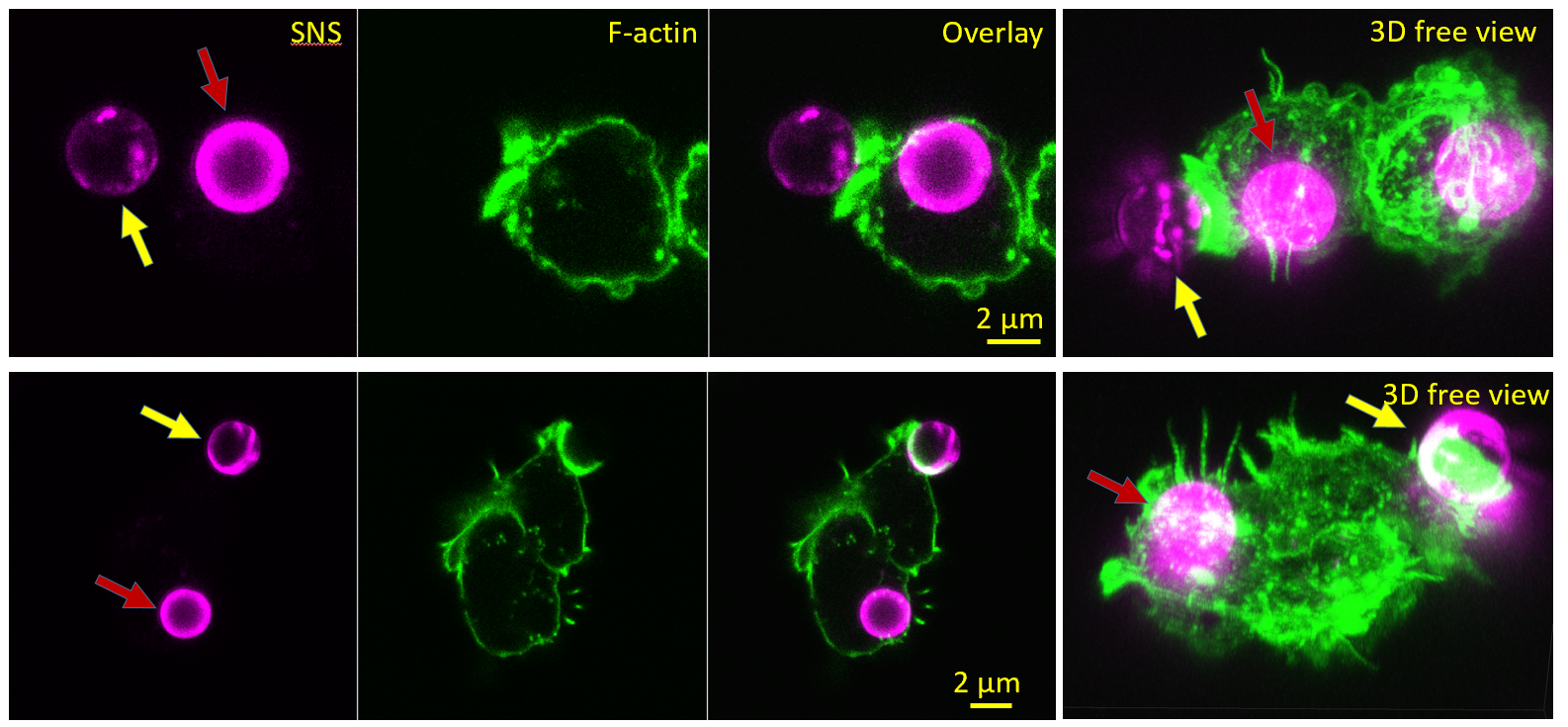

**Fig. S8. The PC exhibits DNase activity over free microbeads (not surface-immobilized).**

Streptavidin-functionalized microbeads (3 µm) were coated with SNS via the avidin-biotin interaction and then added to surface-adherent RAW macrophages. After 30 min of incubation, many microbeads (red arrows) had been internalized by the macrophages and exhibited strong fluorescence, indicating DNase activity within the phagosomes. Several microbeads appeared to be undergoing phagocytosis which is not yet completed and were colocalized with phagocytic cups (PCs), as indicated by F-actin-rich cup structures. Notably, these microbeads (yellow arrows) displayed non-uniform fluorescence across their surfaces, suggesting that phagocytic cups already have DNase activity in response to free microparticles before phagocytic cup closure.

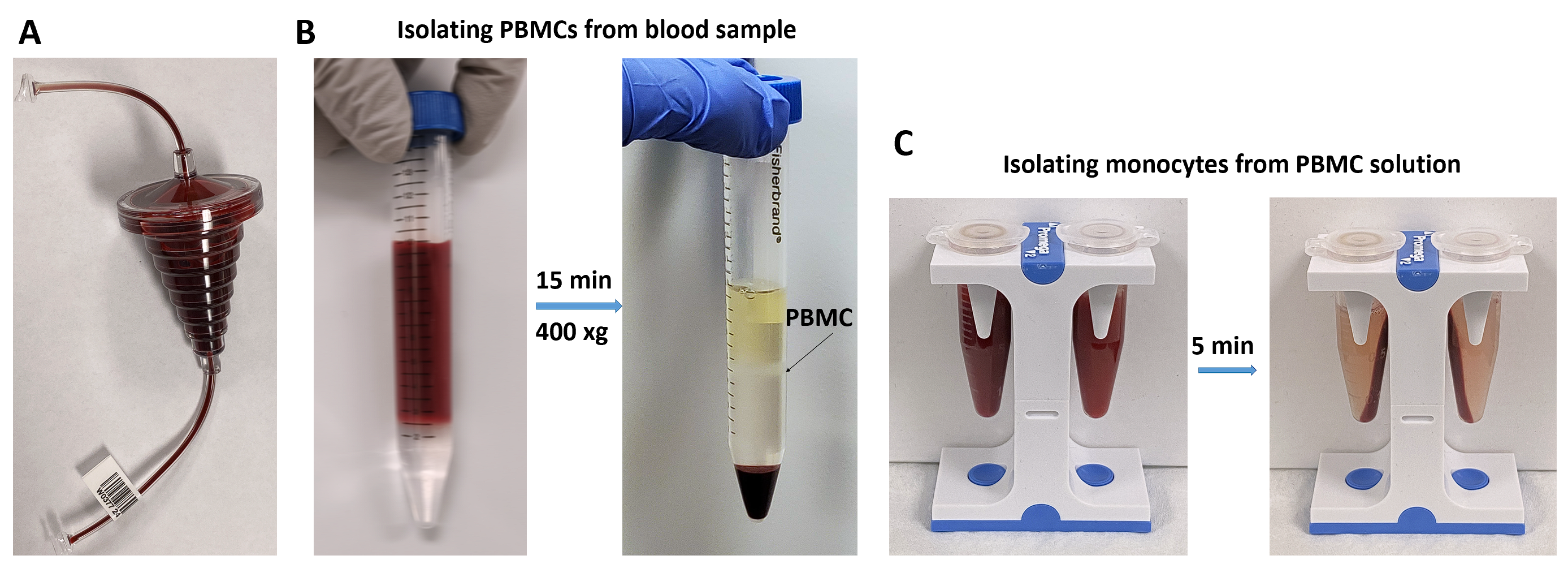

Fig. S9. Human monocyte extraction from white blood cell (WBC)-enriched blood sample

(A) A leukocyte reduction system (LRS) chamber used during blood donation was collected. The chamber contains white blood cell (WBC)-enriched blood sample, providing a source for extracting human monocytes. The blood sample was drained into a 50 mL tube. Phosphate-buffered saline (PBS) supplemented with 2 mM EDTA and 2% fetal bovine serum (FBS) was added to the blood sample at a volumetric ratio of 3:1 (PBS:blood).

(B) 6 mL diluted blood sample was loaded on top of 3 mL gradient medium (the colorless transparent liquid at the bottom of the tube) carefully, forming a clear line between two liquids. The sample was centrifuged at 400 ×g for 15 min. Consequently, red blood cells were spun to the bottom and peripheral blood mononuclear cells (PBMCs) were concentrated between the plasma (the pinkish top layer of liquids) and the gradient medium. The PBMCs were collected by pipetting, and suspended in 2 mL PBS supplemented with 2 mM EDTA and 2% FBS.

(C) Monocytes in the PBMC solution were isolated using a negative selection kit. This kit employs antibody-coated paramagnetic beads to capture all non-monocyte cells, leaving monocytes in the solution. The purified monocytes were subsequently centrifuged and suspended in culture medium for cell culture and experimentation, or alternatively, suspended in a solution of 20% DMSO and 80% FBS for cryogenic preservation.

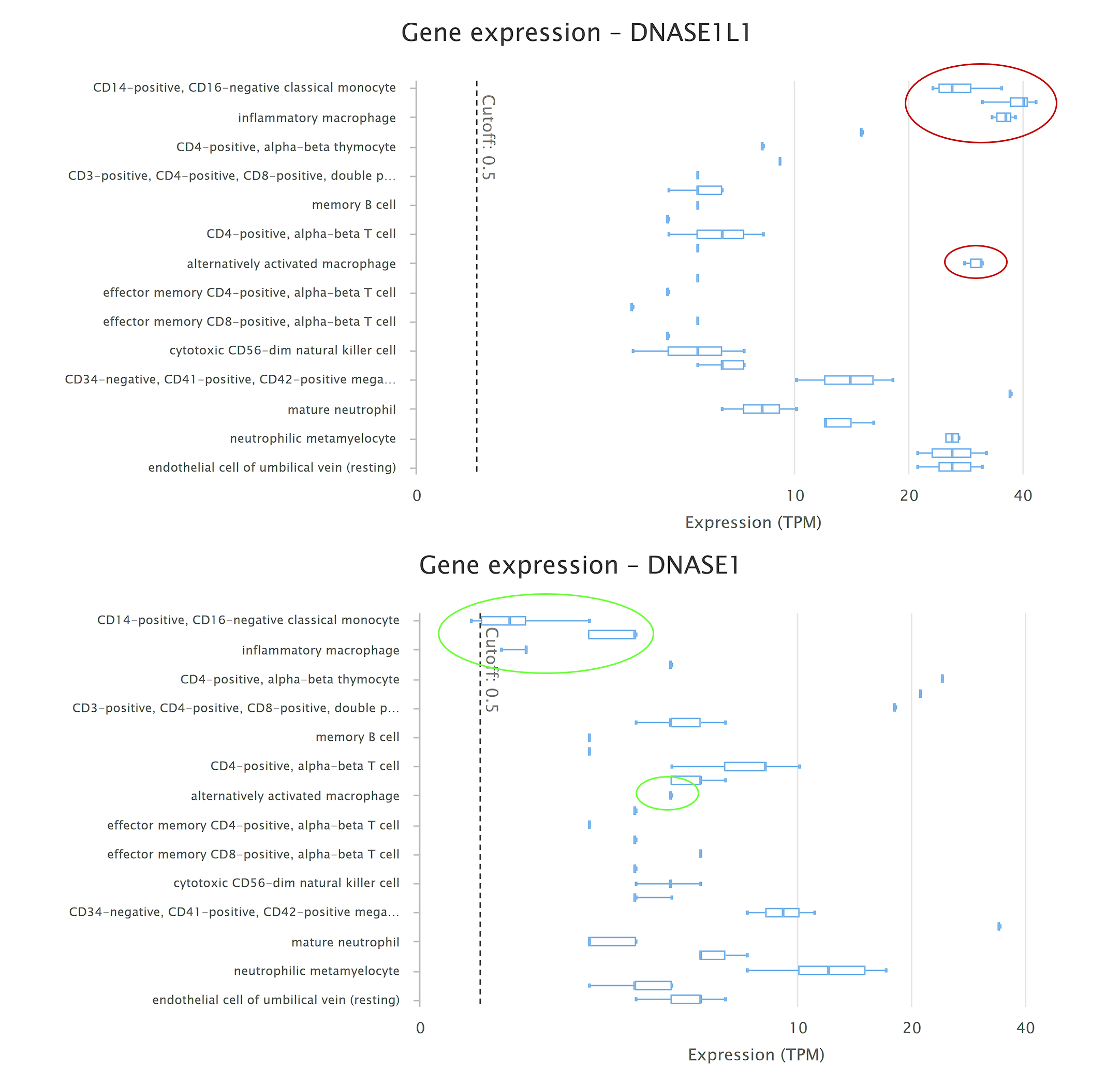

**Fig. S10. Gene expression of DNaseX (a.k.a. DNase I-like 1) in human hematological cells**

The gene expression levels of DNaseX in human hematological cells are available in the Expression Atlas, a comprehensive database maintained by the European Bioinformatics Institute (EBI) that catalogs gene expression across various biological conditions and tissues. According to the data, mRNA of DNaseX is expressed at higher levels in macrophages and monocytes compared to other hematological cells, whereas mRNA of DNase I shows lower expression levels in macrophages relative to other hematological cell types.

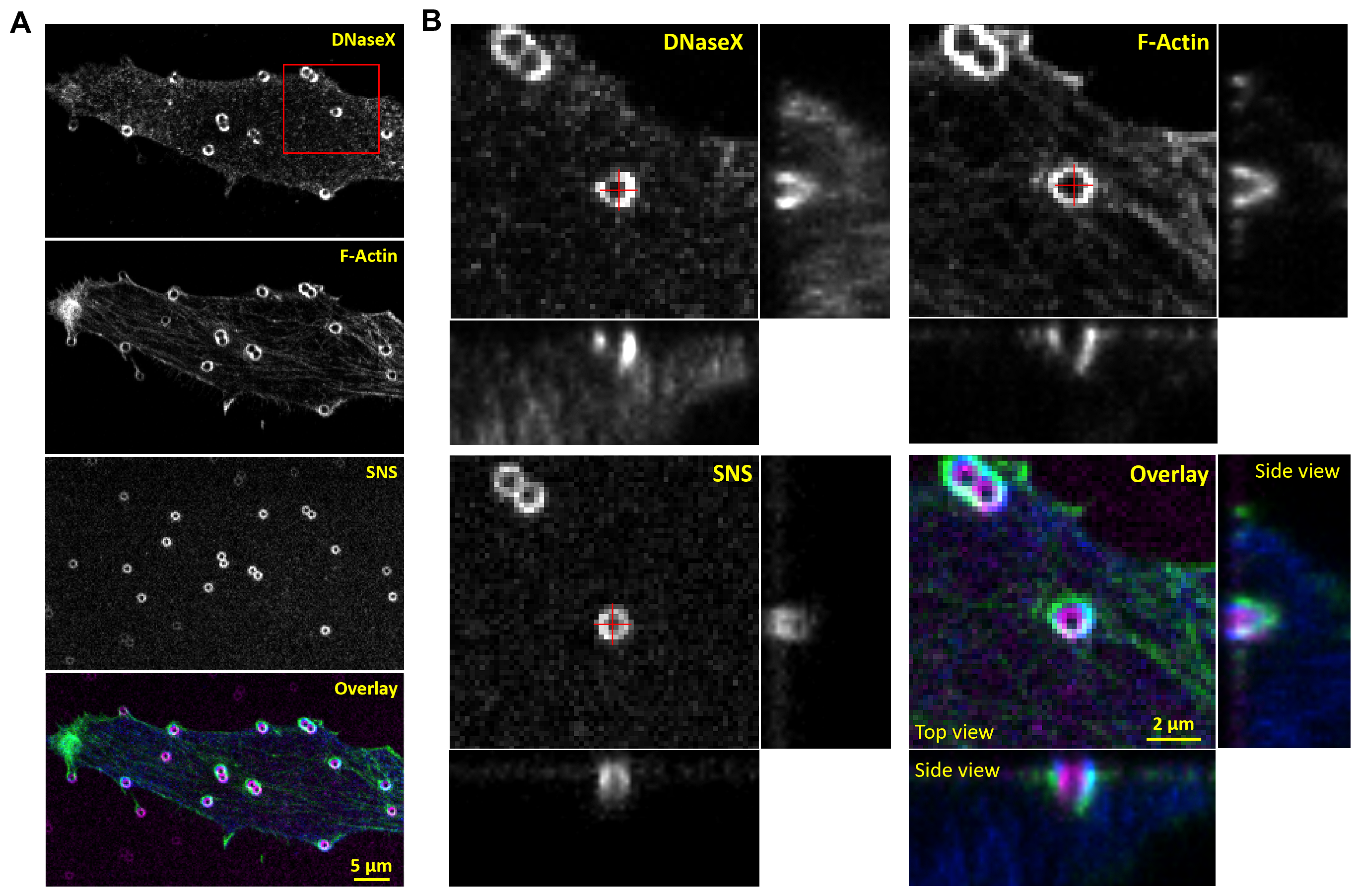

**Fig. S11. DNaseX was recruited onto microbeads beneath the domes of PCs.**

(**A**) Co-imaging of DNaseX (immunostained), F-actin and SNS in a THP-1 macrophage.

(**B**) 3D scan of DNaseX, F-actin and SNS using confocal microscopy. All the three signals are shown to be on the microbeads beneath the domes of the phagocytic cups (PC), not on the substrate regions around microbeads.

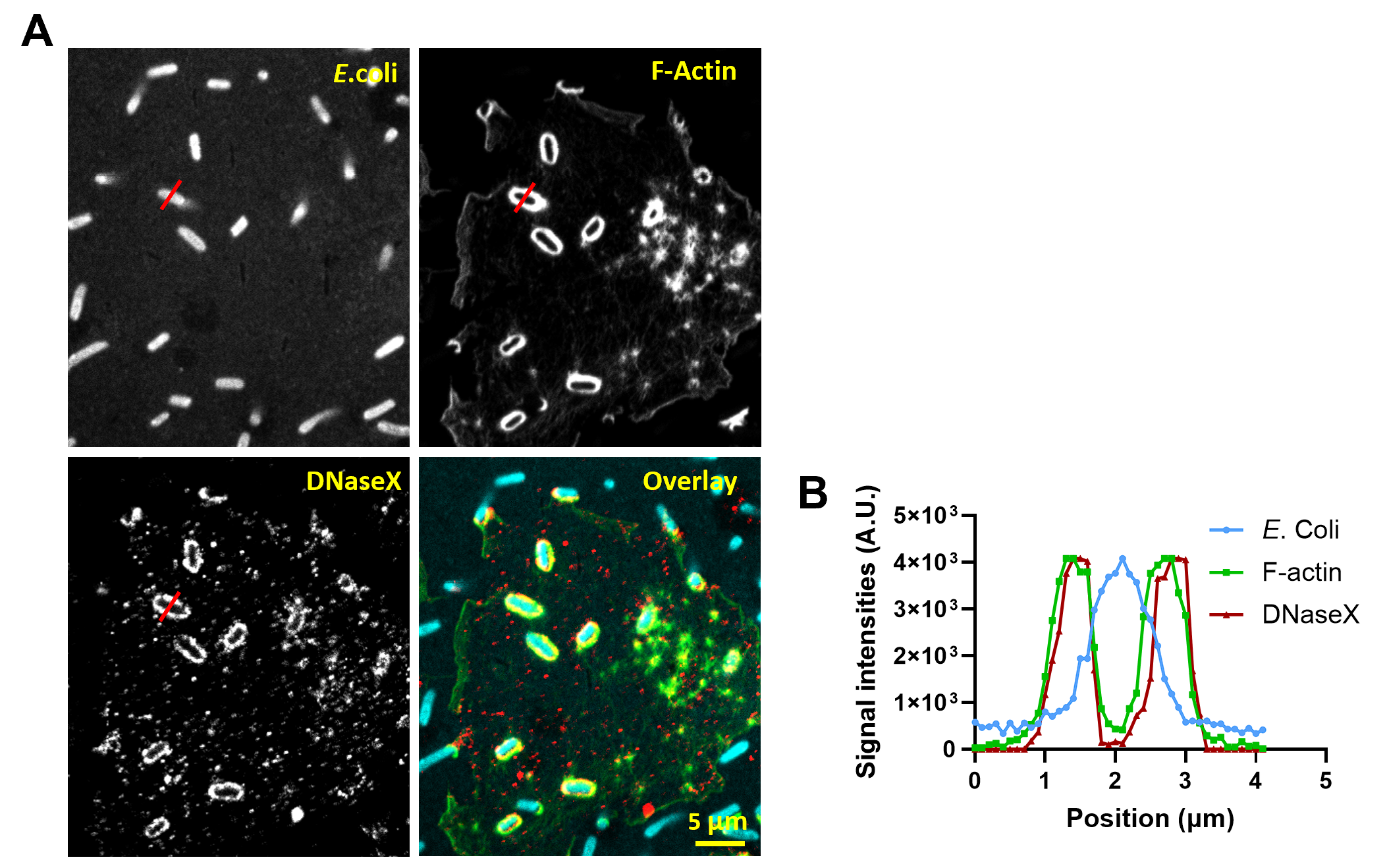

**Fig. S12. DNaseX was recruited to the PCs in response to surface-immobilized *E. coli*.**

(**A**) Co-imaging of *E. coli*, F-actin and immunostained DNaseX underneath a THP-1 macrophage.

(**B**) Line profiles of *E. coli*, F-actin and immunostained DNaseX. The region of analysis is marked with a red line in (A).

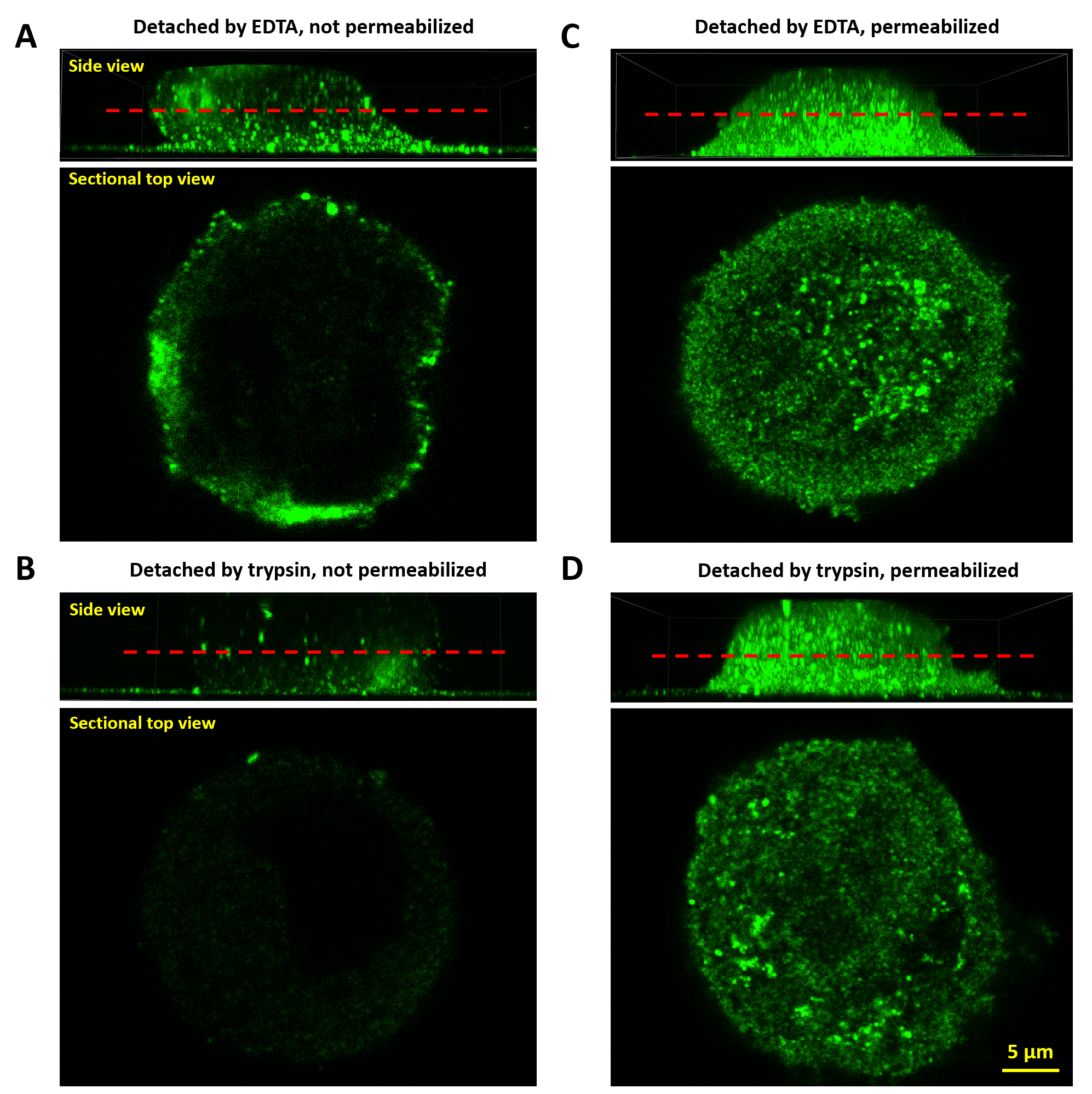

**Fig. S13. DNaseX are found in both plasma membrane and inside cells**

(**A**) Confocal scan of DNaseX on the plasma membrane of macrophages, which were detached by EDTA and not permeabilized during cell fixation. The red dash line indicates the section of 3D imaging displayed in the top view.

(**B**) Confocal scan of DNaseX on the plasma membrane of macrophages, which were detached by trypsin and not permeabilized during cell fixation.

(**C**) Confocal scan of DNaseX on the plasma membrane and in the cytoplasm of macrophages, which were detached by EDTA and permeabilized during cell fixation.

(**D**) Confocal scan of DNaseX on the plasma membrane and in the cytoplasm of macrophages, which were detached by trypsin and permeabilized during cell fixation.

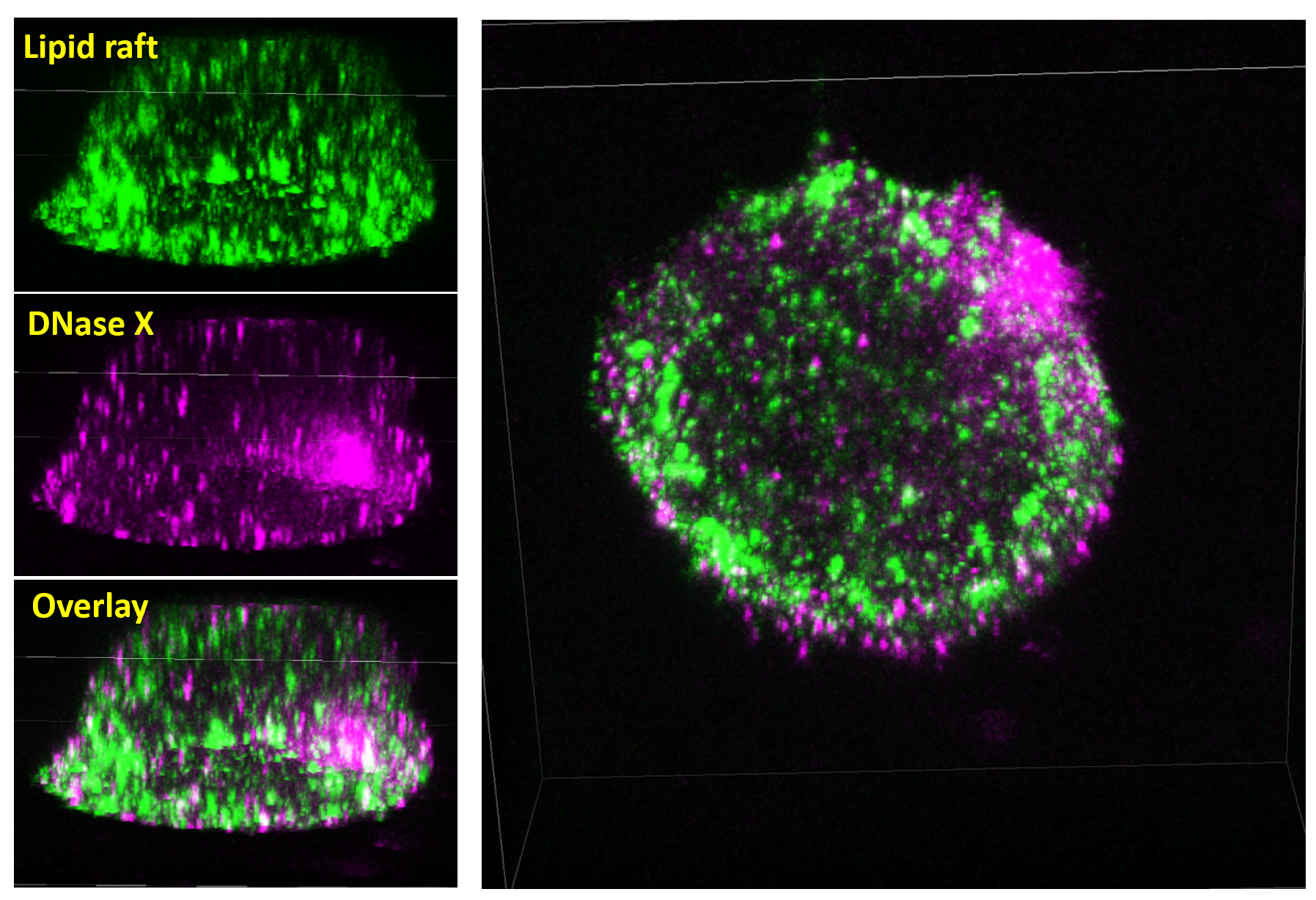

**Fig. S14. No co-localization between DNaseX and lipid rafts on the cell membrane of a macrophage.**

Confocal scanning reconstructed the three-dimensional distribution of lipid rafts and DNaseX clusters on the plasma membrane of a macrophage. No apparent co-localization was observed between these two signals. The lipid rafts were stained with a lipid raft labeling kit (V-34405, Molecular Probes). Related to Video 2.
